# Oral oxycodone self-administration leads to features of opioid addiction in male and female mice

**DOI:** 10.1101/2022.07.19.500655

**Authors:** Richard A. Slivicki, Tom Earnest, Yu-Hsuan Chang, Rajesh Pareta, Eric Casey, Jun-Nan Li, Jessica Tooley, Kavitha Abiraman, Yvan M. Vachez, Drew K. Wolfe, Jason T. Sackey, Robert W. Gereau, Bryan A. Copits, Alexxai V. Kravitz, Meaghan C. Creed

**Affiliations:** Washington University Pain Center, Dept. of Anesthesiology, Washington University in St. Louis; Department of Psychiatry, Washington University in St. Louis; Department of Neuroscience, Washington University in St. Louis; Department of Biomedical Engineering, Washington University in St. Louis

## Abstract

Use of prescription opioids, particularly oxycodone is an initiating factor driving the current opioid epidemic. There are several challenges with modeling oxycodone abuse. First, prescription opioids including oxycodone are orally self-administered and have different pharmacokinetics and dynamics than morphine or fentanyl which have been more commonly used in rodent research. This oral route of administration determines the pharmacokinetic profile, which then influences the establishment of drug-reinforcement associations in animals. Moreover, the pattern of intake and the environment in which addictive drugs are self-administered are critical determinants of the levels of drug intake, of behavioral sensitization, and of propensity to relapse behavior. These are all important considerations when modeling prescription opioid use, which is characterized by continuous drug access in familiar environments. Thus, to model features of prescription opioid use and the transition to abuse, we designed an oral, homecage-based oxycodone self-administration paradigm. Mice voluntarily self-administer oxycodone in this paradigm without any taste modification such as sweeteners, and the majority exhibit preference for oxycodone, escalation of intake, physical signs of dependence, and reinstatement of seeking after withdrawal. In addition, a subset of animals demonstrate drug taking that is resistant to aversive consequences. This model is therefore translationally relevant and useful for studying the neurobiological substrates of prescription opioid abuse.

## Introduction

Misuse of prescription opioids continues to present a significant public health and economic burden worldwide. In 2019 in the United States, 9.7 million people reported misuse of prescription pain relievers, and one-third of this population reported misusing of oxycodone [1]. Despite its importance, modeling prescription opioid abuse in rodents has been challenging [2-4]. Certain aspects of prescription opioid use, such as the oral route of administration and regular, long-term intake, may contribute to abuse liability and propensity toward relapse. As such, a representative preclinical model of prescription opioid abuse is critical for understanding the neurobiology underlying this disorder.

In contrast to many reinforcing substances, prescription opioids such as oxycodone (OxyContin ™, Tylox™ or Percodan™, abbreviated OXY) are most often orally self-administered: among chronic and recreational opioid users, oral self-administration is decidedly preferred over non-oral routes (e.g., insufflation, injection) [5]. The pharmacokinetic and pharmacodynamic profiles of opioids and therefore their physiology, time-course of action, and metabolism vary widely across routes of administration [6] and among subclasses of opioids [7-9]. Specifically, OXY exhibits higher bioavailability in the GI tract relative to morphine or fentanyl when administered orally [7-11]. It is also important to note that while all prescription opioids are thought to exert their reinforcing properties via activation of µ-opioid receptors, OXY exhibits differentially binding affinities to other subclasses of opioid receptors relative to morphine or fentanyl [12]. Further, the OXY is subject to different primary enzymatic degradation pathways relative to other µ-opioid agonists [13]. These differences between classes of µ-opioid agonists with respect to receptor binding, signaling and enzymatic degradation result in differential engagement of downstream signaling pathways with variable efficacy and rates of receptor desensitization [14], Emery 2016. The rate at which brain levels of a drug rise is critical for establishing associations between drug intake and its euphoric properties [15, 16]. Since OXY exhibits slower absorption and distribution orally relative to intravenous or subcutaneous routes [17] modelling oral OXY intake would provide a potentially more relevant comparsion to the clnical features of OXY intake and misuse.

The pattern of intake of drugs of abuse is also relevant for establishing learned drug associations [18], which contribute to the subjective reinforcing properties of the drug, escalation of intake, and propensity toward reinstatement [19-25]. Specifically, intermittent drug access, which produces fluctuating brain drug levels, exacerbates mesolimbic adaptations induced by both cocaine [26, 27] and opioid self-administration [28, 29] and increases motivation to self-administer the drug later on [30]. In contrast, prescription opioids, particularly extended release formulations, are designed to deliver a consistent analgesic response without rapid fluctuations in drug concentrations in the brain. Recapitulating this continuous access pattern should also be considered when modeling prescription opioid abuse. Prescription opioids are also most frequently self-administered in a familiar setting such as the home (see Caprioli et al. (2007) for a review[31]). Environmental context (i.e., setting of drug taking) plays an important role in both drug taking and reinstatement, a finding that has been well-documented in humans and recapitulated by animal models [31-35]. Yet, prior models of oral opioid self-administration have taken place in novel contexts (e.g., operant chamber) and under food restriction [2-4]. Thus, modelling continuous prescription opioid access in a familiar environment may provide a more clinically relevant comparison to prescription opioid self-administration

Previous investigations have investigated the oral route of self-administration by implementing two-bottle choice procedures, where animals are free to choose between bottles containing drug- and drug-free solutions. Because the opioid alkaloid structure confers a bitter taste to opium derivatives, sweeteners were added to encourage animals to self-administer opioids, or adulterants to the drug-free bottle to control for aversive taste [36-38]. However, the addition of these extraneous reinforcers or adulterants may introduce experimental confounds [39]. For example, sucrose itself can support seeking after abstinence [40], disrupt overall patterns of fluid intake [41], and stimulate dopamine and opioid receptors in the mesolimbic pathway [42-45]. Therefore, we sought to develop a preclinical model for studying prescription opioid use disorder that recapitulates continuous access pattern and oral route of administration without adulterants.

Here, we characterize a home cage-based, oral OXY self-administration paradigm. Our paradigm produced behavioral features of opioid use disorder (OUD) as defined by the fifth version of the diagnostic and statistics manual (DSM)[46] including escalation of drug intake, physical signs of dependence, drug craving after withdrawal and drug use despite negative consequences. These behaviors were also accompanied by neural changes that included potentiation of excitatory synapses in the nucleus accumbens (NAc), which is a conserved neurobiological substrate of drug reinstatement [47-49]. We demonstrate that the high-throughput nature of our model is valuable for studying individual variability in both the amount and pattern of drug intake, as well as individual susceptibility to aversion-resistant drug consumption and reinstatement of drug-seeking behaviors after withdrawal.

## Materials and Methods

### Subjects

A total of 112 adult mice (C57BL/6J background; age, 8–16 weeks) were used in this study. Both males (n=58) and females (n=67) were included and balanced across experiments and treatment groups. Mice were housed in standard mouse vivarium caging and kept on a 12 h light/dark cycle (light onset, 7:00 A.M.; light offset, 7:00 P.M., temperature (22-26°C), in-cage humidity (22-50%). Mice had ad libitum access to food and were single housed during self-administration experiments. Experiments were run in successive waves of treatment- and sex-matched cohorts with up to 20 mice being run simultaneously. All studies were approved by the internal animal care and use committee at Washington University in St. Louis.

### OXY self-administration

<u>Habituation:</u> Mice were single housed and cages were equipped with an in-cage two-bottle choice apparatus prior to the start of the escalation protocol, during which two 15 mL conical tubes equipped with a drinking valve in a 3D printed in-cage holder was the only source of drinking water (water was supplied in both tubes). <u>Phase I:</u> Mice in the experimental group (“OXY mice”) were supplied with a single bottle of oxycodone hydrochloride (Sigma Aldrich) dissolved in drinking water as their only source of liquid; 0.1 mg/mL was available for 24 hours, 0.3 mg/mL was available for 48 hours, and 0.5 mg/mL was available for 48 hours (Fig 1A), based on an operant protocol developed by Phillips et al. [2]. Mice in the control group (“CTRL mice”) had access to drinking water only throughout the protocol. The position of the bottle within the two-bottle apparatus was switched daily. <u>Phase II:</u> OXY (1.0 mg/mL in drinking water) and drinking water were supplied to OXY mice *ad libitum* in the two-bottle choice apparatus, position of bottles was balanced within groups at the population level. Both solutions were available 24 hours a day for 7 days. Drinking water was supplied in both bottles for CTRL mice. <u>Withdrawal:</u> The two-bottle lickometer was removed from the home cage after the 7-day two-bottle choice period, and standard cage-top water bottles were replaced as the source of drinking water. Separate groups of mice were used for the responder analysis, OXY seeking, quinine adulteration and electrophysiology studies outlined below.

**Figure 1.**
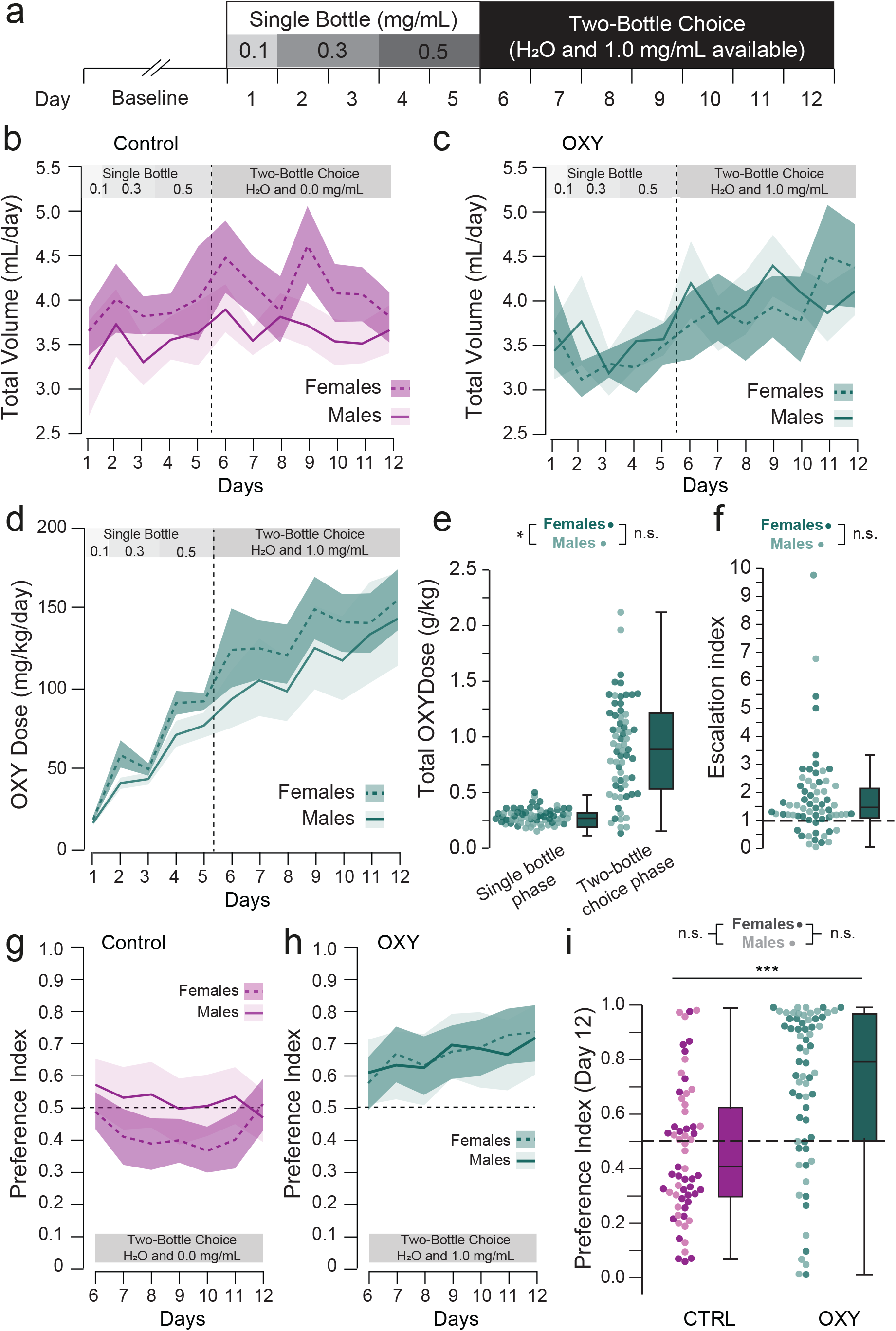
Mice voluntarily consume oxycodone in a two-bottle choice paradigm. **(a)** Schematic of experimental paradigm (n = CTRL: 33F/28M, OXY: 35F/30M). **(b-c)** Total daily volume of liquid consumed throughout the self-administration protocol for **(b)** control and **(c)** OXY-administering mice. Mice with OXY available consumed comparable total cumulative volumes of liquid (F_Drug_ = 0.07, p=0.224); although there was a significant interaction between drug and day (F_Drug•Day_ = 2.47, p < 0.005). **(d)** Daily self-administered doses of OXY. Self-administered OXY doses increased with concentration throughout the protocol (F_Day_ = 52.76, p<0.001), during which female mice self-administered higher daily doses than males during the single-bottle phase (F_sex_ = 12.81, p < 0.001) but not two-bottle choice phase (F_Sex_ = 1.51, p = 0.224) **(e). (f)** No difference in escalation index (Dose_Day12_ / Dose_Day6_) was observed between male and female mice (F_sex_ = 0.69, p = 0.409). **(g-h)** Preference for the OXY-containing bottle over the 7 days of the two-bottle choice phase for **(g)** control and **(h)** OXY-administering mice. OXY mice only exhibited a preference for the OXY-containing bottle (F_drug_ = 17.682, p<0.0001) which increased over successive days (F_Day•Drug_ = 3.78, p = 0.001 with no sex differences (F_sex_ = 1.636, p=0.13). **(i)**There was no systematic preference for drinking water from the left or right bottle in CTRL mice, while OXY mice consumed significantly more OXY solution relative to drinking water when available (Preference Index = OXY Volume _(Day 12)_/Total Volume_(Day 12)_, F_drug_= 22.67, p<0.001). There were no sex differences in OXY preference (F_sex_ = 0.188, p = 0.665, F_sex•drug_ = 1.207, p = 0.274) ***p<0.001.

### Home cage lickometer devices

Initial iterations of the OXY two-bottle choice procedure did not include lickometer devices, and only focused on volumetric analysis. Devices used in these experiments are low-cost and open-source, with detailed parts lists, code and fabrication instructions published and available on-line [50, 51]. Drinking water and OXY solutions were provided using an in-cage two-bottle lickometer apparatus which detects interactions with two separate drinking spouts via a pair of photobeams [51]. In a subset of experiments, cages were also equipped with a wireless passive infrared (PIR) activity monitor [50] to measure mouse activity levels (Models 4430 and 4610 from MCCI Corporation, Ithaca NY). Lickometer counts on each lickometer and PIR activity data were transmitted via Low frequency Radio Wide Area Network (LoRaWAN) using Internet of Things (IoT) infrastructure (The Things Network, Netherlands), saved in a cloud-database (InfluxDB), and visualized with an online dashboard (Grafana). The wireless lickometer and PIR sensors were also equipped with back-up microSD cards and liquid volumes were measured manually each day.

### Responder analysis, Lickometer preference index and circadian index

Responder analysis was performed on a subset of animals in which lickometer and volumetric data were available. Volumetric intake was used to calculate whether animals were high or non-responders. Animals were considered high responders if the overall amount of OXY consumed relative to water was over >90%, and non-responders if the OXY solution accounted for less than 50% of their total overall volume consumed over the 7 day 2 bottle choice period. The Lickometer Preference Index ([Counts on OXY-containing bottle] / [Total Counts], Fig 2A), *was* used to determine the number of general interactions between water and OXY containing bottles and the Circadian Index ([Total Counts during the dark cycle] / [Total Counts during the light cycle]; Fig 2B) was used to evaluate the microstructure of drinking activity as a proxy for circadian behavior.

**Figure 2.**
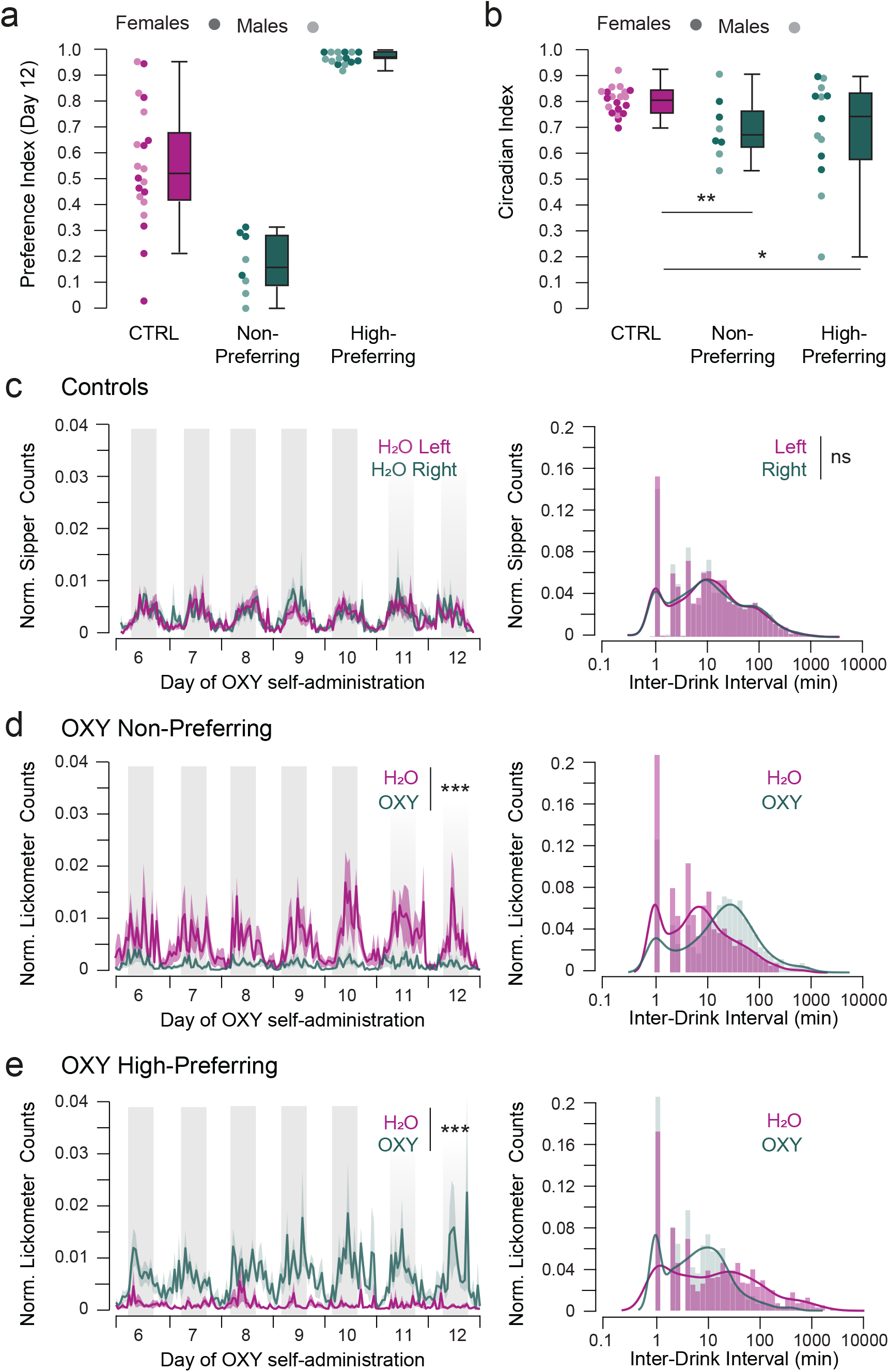
Non-preferring and high-preferring mice exhibit circadian disruption and distinct patterns of drug intake. **(a)** Preference index of CTRL (11F/10M, 0.55 ± 0.05), Non-Preferring (4F/4M, 0.17 ± 0.04) and High-Preferring (7F, 6M, 0.98 ± 0.01) OXY mice with lickometer data represented. **(b)** Circadian index was altered in both Non- (0.70 ± 0.04) and High-Preferring mice (0.68 ± 0.06), relative to controls (CTRL: 0.80 ± 0.01, F_Group_ = 3.42, p = 0.044). **(c-e)** Control mice exhibited no systematic difference in lickometer counts (t=0.29, p=0.77) or inter-drink intervals on the left vs right bottle (KS: 0.07, p = 0.994). In contrast, Non-preferring and High-preferring mice exhibited proportionately more counts on the H_2_O-containing (t=11.93, p<0.001) and OXY-containing (t=12.84, p<0.001) bottles. Similarly, the distribution of interdrink-intervals on the OXY-paired bottle was significantly different from the distribution of CTRL mice for both Non-preferring (KS: 1.36, p = 0.049) and High-preferring mice (KS: 1.62, p=0.011). *p<0.05, **p<0.01, ***p<0.001.

### OXY Seeking Test

In a subset of mice (n_CTRL_ =9M/10F, n_OXY_ = 13M/13F), an OXY seeking test was performed under extinction conditions. 21 days following removal of the lickometer device, a lickometer device containing 2 empty bottles was re-introduced to the cage for a 60-minute probe trial, and interactions with the lickometer tubes as detected by the photobeam were recorded. Trials were run between 12pm and 2:30 pm for all mice.

### Quinine Adulteration

In a subset of mice (n_CTRL_ = 8M/7F, n_OXY_ = 8M/7F) we tested resistance of OXY drinking to quinine adulteration. Following the conclusion of the 7 days of two-bottle choice protocol, increasing concentrations of quinine were added to the OXY-containing bottle (OXY mice), such that OXY mice had the choice between standard drinking water (one bottle) and 1.0 mg/mL OXY + quinine. CTRL mice had the choice between standard drinking water (one bottle) and water + quinine. Mice were given 48 hours at each quinine concentration in increasing order: 125 µM, 250 µM, 325 µM, volumes consumed from each tube were recorded daily.

### Physical Withdrawal Signs

Markerless pose estimation was used to quantify physical signs of opioid dependence in a subset of mice (n_CTRL_ = 6M/6F, n_OXY_ = 8M/12F) <u>Acquisition:</u> After either 24 hours (acute) or 21 days (protracted) of withdrawal, mice were habituated to the testing room for at least 1 hour. Fifteen minute videos were acquired of individual mice at > 60 fps in a 4” x 4” x 8” plexiglass chamber with an unobstructed background and uniform overhead lighting. Videos were scored for signs of withdrawal (jumps, tremors, movement speed) and anxiety related behaviors (grooming, rearing, climbing). Markerless pose estimation was conducted using DeepLabCut™ (DLC, version 2.2b6 [46,47]): 12 frames each from 64 videos representing all treatment conditions were extracted using k-means clustering and subsequently manually annotated for the following 9 body parts: left ear, right ear, left forepaw, right forepaw, left hind paw, right hind paw, snout, tail base, back. The training fraction was set to 0.95, and the resnet_50 network was trained for 1,030,000 iterations. A train error of 1.82 and test error of 11.47 were achieved with a cutoff value of p=0.6. DLC tracking data and generated videos were then imported to the Simple Behavioral Analysis (SimBA) project workflow (version 1.2 [52]). Within SimBA, single animal, 9-body part supplied configuration was used to extract behavioral features from the pose estimation data after outlier and movement correction (both parameters set to 7x outside the interaural distance. Extracted frames from four independent videos were annotated to build classifiers for the following behaviors: “climbing”, “jumping”, “tremor”, “rearing, and “grooming”. Individual models were trained using a random forest machine model with 2000 estimators, and a training fraction of 0.2 (default hyperparameters). Videos were analyzed using the random forest model, with the following p-cutoffs and minimum behavioral bout lengths for each of the following behaviors: “climbing (p = 0.26, 35 ms)”, “jumping (p = 0.4, 35 ms)”, “tremor (p=0.0495, 50 ms”, “rearing (p = 0.45, 70 ms)”, and “grooming (p = 0.38, 70 ms)”, wet dog shakes were manually scored by an investigator blind to treatment condition. The SimBA pipeline is built primarily on scikit-learn [53], OpenCV[54], FFmpeg45[55], and imblearn[56]. Number of events are reported for jumps, wet dog shakes and tremor episodes, whereas rearing, grooming and climbing are reported in time spent exhibiting each behavior. Euclidean distance of displacement of all body parts was also extracted from the pose-estimation data and averaged to achieve one ‘movement’ score for each subject.

### Patch Clamp Electrophysiology

Coronal mouse brain slices, 220 μm in thickness were prepared in cooled artificial cerebrospinal fluid containing (in mM): 119 NaCl, 2.5 KCl, 1.3 MgCl, 2.5 CaCl_2_, 1.0 Na_2_HPO_4_, NaHCO_3_ 26.2 and glucose 11, bubbled with 95% O_2_ and 5% CO_2_. Slices were kept at 30-32°C in a recording chamber perfused with 2.5 mL/min artificial cerebrospinal fluid. Visualized whole-cell voltage-clamp recording techniques were used to measure spontaneous and synaptic responses of NAc shell MSNs. Holding potential was maintained at -70 mV, and access resistance was monitored by a depolarizing step of -10 mV each trial, every 10 s. Currents were amplified, filtered at 2 kHz and digitized at 10 kHz. All experiments were performed in the presence of picrotoxin (100 μM) to isolate excitatory transmission. Internal solution contained (in mM): 130 CsCl, 4 NaCl, 5 creatine phosphate, 2 MgCl_2_, 2 Na_2_ATP, 0.6 Na_3_ GTP, 1.1 EGTA, 5 HEPES and 0.1 mm spermine. Synaptic currents were electrically evoked by delivering stimuli (50 – 100 μs) every 10 seconds through bipolar stainless-steel electrodes. The AMPAR component was calculated as the peak amplitude at -70 mV, The NMDAR component was estimated as the amplitude of the outward current at +40 mV after decay of the AMPA current, measured 50 msec after the electrical stimuli (Fig 5B). Paired-pulse ratio PPR was calculated as the ratio of the second to first baseline-subtracted peak elicited with an ISI of 50 msec (Fig 5B).

### Statistical Analyses

Photobeam break data on the lickometer devices was extracted and analyzed using custom python code (SipperViz graphical user interface, https://github.com/earnestt1234/SipperViz). Statistics were performed in python (3.7 using Spyder 4.1.5), using the pinguoin (0.3.10) and statsmodels (0.10.0) packages. Data were analyzed with repeated measures ANOVA or two-factor ANOVA where appropriate, followed by post-hoc t-tests. Sex and Treatment group were included as between-subject factors for all analyses. Individual data points alongside boxplots marking interquartile distributions are shown; aggregate data are presented as means +/- 90% confidence intervals.

## Results

### Mice exhibit preference for OXY over drinking water and escalate OXY intake over time

We mimicked a prescribed course of OXY by escalating concentrations of OXY in the drinking water in a single-bottle phase, followed by a two-bottle choice phase where mice could choose between OXY solution (1.0 mg/mL in drinking water) and unadulterated drinking water (Fig 1A). During the single bottle phase, mice increased their daily OXY consumption (Fig 1B-D), which was predicted due to increasing concentrations of OXY in their drinking water (0.1, 0.3, 0.5 mg/mL). However, mice continued to escalate their drug intake during the two-bottle choice phase where the drug concentration remained constant (1.0 mg/mL) and drinking water was also available ad libitum (Fig 1D-F). While body weight was systematically higher in males relative to females, it was not significantly different between control and OXY mice (CTRL male = 27.44 ± 0.55, female = 21.29 ±0.54, OXY male = 25.65 ± 0.56, female = 21.19 ± 0.39, F_sex_ = 88.71, p< 0.0001, F_drug_ = 2.71, p = 0.102, F_sex*drug_ = 2.153, p = 0.145).

During the two-bottle choice phase, mice also exhibited a significant *preference* for 1.0 mg/mL OXY over drinking water at the population level (Fig 1G-I), despite the lack of sweeteners or tastants in the drinking water or OXY solution. There was no systematic preference for the left or right bottle in CTRL mice (Fig 1G,I), while OXY mice exhibited higher levels of responding on the OXY-containing bottle (Fig 1H,I), with no significant sex differences with respect to the total amount of OXY consumed, escalation, or preference for OXY. However, female mice consumed significantly higher weight-adjusted volumes of OXY during the single-bottle phase (Fig 1C,D, Supp Fig 1A-C).

### Mice exhibit individual differences in OXY intake

There was considerable variability in OXY preference among individual mice; while no mice completely avoided OXY, a subset did prefer unadulterated drinking water over the OXY solution (Fig 1I, Supp Fig 1D-G). We therefore evaluated a subset of mice for variability in OXY preference, and how OXY preference would alter the microstructure and circadian pattern of drinking behavior (n=11F, 10M CTRL, and n=11F/12M OXY). We observed that in a subset of mice (n=4F/4M), OXY accounted for less than 50% of the total liquid intake. We define these mice as <u>OXY Non-preferring</u>. In contrast, we observed a larger number of mice that consumed over 90% of their total fluid intake from the OXY solution (Fig 1I, Supp Fig 1D-G), which we refer to as <u>OXY High-preferring</u> (n=7F/8M).

### OXY intake alters the microstructure and circadian pattern of drinking behavior regardless of individual preference

In addition to total volumes, the home-cage lickometer device we employed registered interactions with each bottle, which allowed for examination of patterns of drug intake. There was no systematic preference for either the left or right H_2_O-containing bottle in control mice (n=11F/10M, Fig 2A), and no difference in the distribution of inter-drink intervals on the left or right bottles (Fig 2C) in control mice. The circadian index was significantly altered in both Non-preferring and High-OXY preferring mice relative to controls (Fig 2B), which was driven by an increase in both water and OXY interactions during the light cycle in OXY-consuming mice (Fig S2A-F. As expected, OXY Non-preferring mice avoided the OXY-containing bottle and had shorter inter-drink intervals on the water-containing bottle relative to the OXY-containing bottle (Fig 2D). By contrast, OXY High-Preferring mice exhibited significantly higher lickometer counts on the OXY-containing bottle and shorter inter-drink intervals on the OXY-containing bottle (Fig 2E). These patterns in intake microstructure were recapitulated at the individual subject level in male and female mice (Supp Fig 3A-F).

### OXY self-administration induces physical signs of intoxication and withdrawal

Acute injections of OXY [57-59], or vapor exposure [60] to opioids induce hyperactivity, which are thought to reflect increased dopamine release in the NAc [61-63]. To examine this, we equipped our home cages with PIR sensors to detect locomotor activity during the self-administration protocol (Fig 3A,B). OXY mice exhibited higher levels of activity than CTRL mice during both the single- and two-bottle choice phases of self-administration (Fig 3D-E), suggesting that voluntary oral OXY self-administration results in sufficient brain levels to induce hyperactivity [58, 64, 65]. We found no sex differences in OXY-induced locomotor activity (Fig S4A-C).

**Figure 3.**
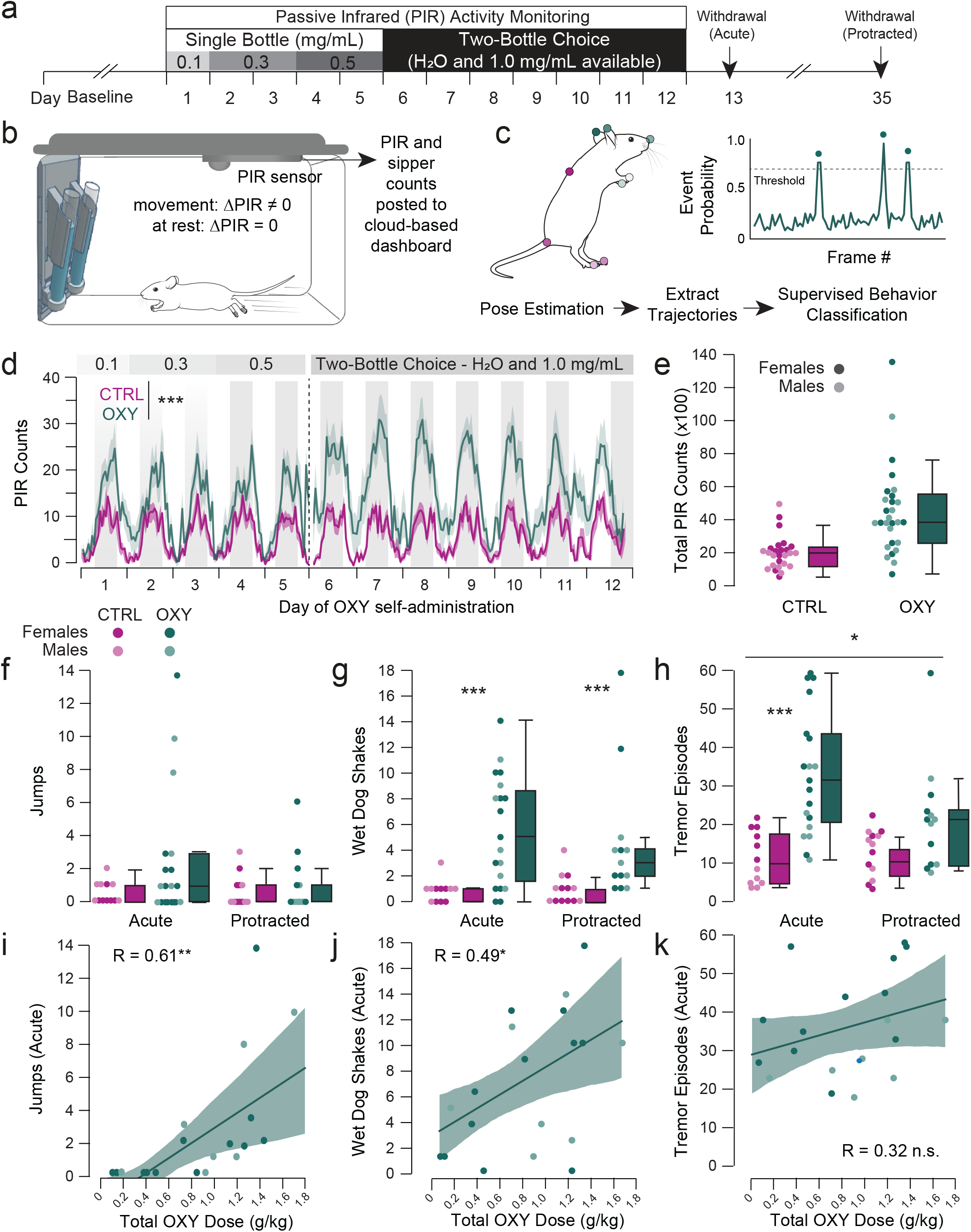
OXY self-administration induces somatic signs of intoxication and withdrawal. **(a)** Experimental schematic. **(b)** Home cages were equipped with PIR sensors to monitor activity during the single- and two-bottle choice phases of OXY-self administration. **(c)** Withdrawal symptoms were quantified after 24 hours and after 3 weeks of withdrawal using post estimation and supervised behavioral classification. **(d-e)** OXY mice exhibited significantly higher homecage activity during self-administration relative to control mice (CTRL: 2072.189 ± 196.455 n=10F/10M, OXY = 4530.370, n=15F/13M, F_sex_ = 0.956, p = 0.333, F_drug_=19.549, p<0.001). **(f)** There was no significant effect of OXY or sex on jumping behavior at the acute (CTRL n=6F/6M: 0.5 ± 0.19, OXY n=12F/8M: 2.37 ± 0.91) or protracted time points (CTRL n=7F/6M: 0.62 ± 0.27, OXY n=7F/7M: 1.0 ± 0.49; F_Drug_ = 3.229, p = 0.079, F_Sex_ = 1.937, p = 0.170, F_TimePoint_= 0.337, p = 0.94). **(g)** There a significant effect of OXY on the number of wet dog shakes mice exhibited at both the early (CTRL: 0.83 ± 0.24, OXY: 5.53 ± 0.97) and protracted time points (CTRL: 0.92 ± 0.33, OXY: 4.54 ± 1.38; F_Drug_ = 17.798, p < 0.001, F_Sex_ = 0.108, p = 0.743, F_TimePoint_ = 0.239, p = 0.627). **(h)** There a significant effect of OXY on the number of tremor episodes exhibited, with OXY mice exhibiting more episodes at the acute (CTRL: 18.25 ± 1.64, OXY: 36.32 ± 3.01) but not protracted time point (CTRL: 16.08 ± 1.59, OXY: 24.08 ± 3.61; F_Drug_= 20.514, p < 0.001, F_Sex_ = 6.657, p = 0.013, F_TimePoint_ = 5.750, p = 0.020). **(i-k)** The number of jumps and wet dog shakes, but not tremor episodes in acute withdrawal was significantly correlated with the amount of self-administered OXY(p=0.004, 0.028, 0.169, respectively). *p<0.05, **p<0.01, ***p<0.001.

We also determined whether withdrawal from oral OXY self-administration induces physical withdrawal signs. During withdrawal from opioids such as morphine or heroin, mice exhibit characteristic withdrawal signs including jumps, wet dog shakes and tremors [66, 67]. Relative to CTRL mice, OXY mice exhibited significantly more jumps (Fig 3F), wet dog shakes (Fig 3G) and paw and body tremor episodes (Fig 3H) in acute opioid withdrawal 24 hours following sipper OXY removal. The increase in wet dog shakes was attenuated but still apparent in OXY mice in protracted (3 weeks) withdrawal (Fig 3H). The number of jumps and wet dog shakes correlated significantly with the total OXY dose an animal consumed over the 12 day protocol (Fig 3I, J), while the number of tremor episodes did not (Fig 3K).

Finally, we quantified grooming and rearing, which have been used as a proxy for anxiety-like behavior and have also been reported to be increased following withdrawal from opioids[68]. OXY mice did not exhibit significantly more rearing or grooming relative to CTRL mice (Supp Fig 5A-C). There were no significant differences between male and female mice for any physical symptom measured. Together, these results demonstrate that oral OXY self-administration induces physical signs of dependence that are consistent with prior reports of an opioid withdrawal syndrome in rodents.

### The two-bottle choice paradigm induces drug seeking under extinction conditions

Drug seeking behavior is operationally defined as performance of an action that previously led to consumption of the drug, in the absence of the drug itself [69, 70]. In the case of intravenous drug self-administration, this response may be operationalized as lever-pressing or nose-poking on an operant device that previously resulted in drug infusions. In the current paradigm, OXY was delivered through a lickometer device which was distinct from the standard cage-top water bottle. Therefore, we assayed drug seeking behavior by measuring photobeam breaks on the lickometer devices under extinction conditions, with no OXY or drinking water in the tubes (Fig 4A). After 3 weeks of forced abstinence, OXY mice exhibited significantly higher lickometer counts in extinction than CTRL mice (Fig 4B-C). This drug seeking behavior was not significantly different between males and females (Fig 4D Supp Fig 6A-F) and did not correlate with the total amount of OXY consumed during the self-administration paradigm (Fig 4E). This establishes that the two-bottle choice paradigm supports drug seeking behavior that persists after abstinence.

**Figure 4.**
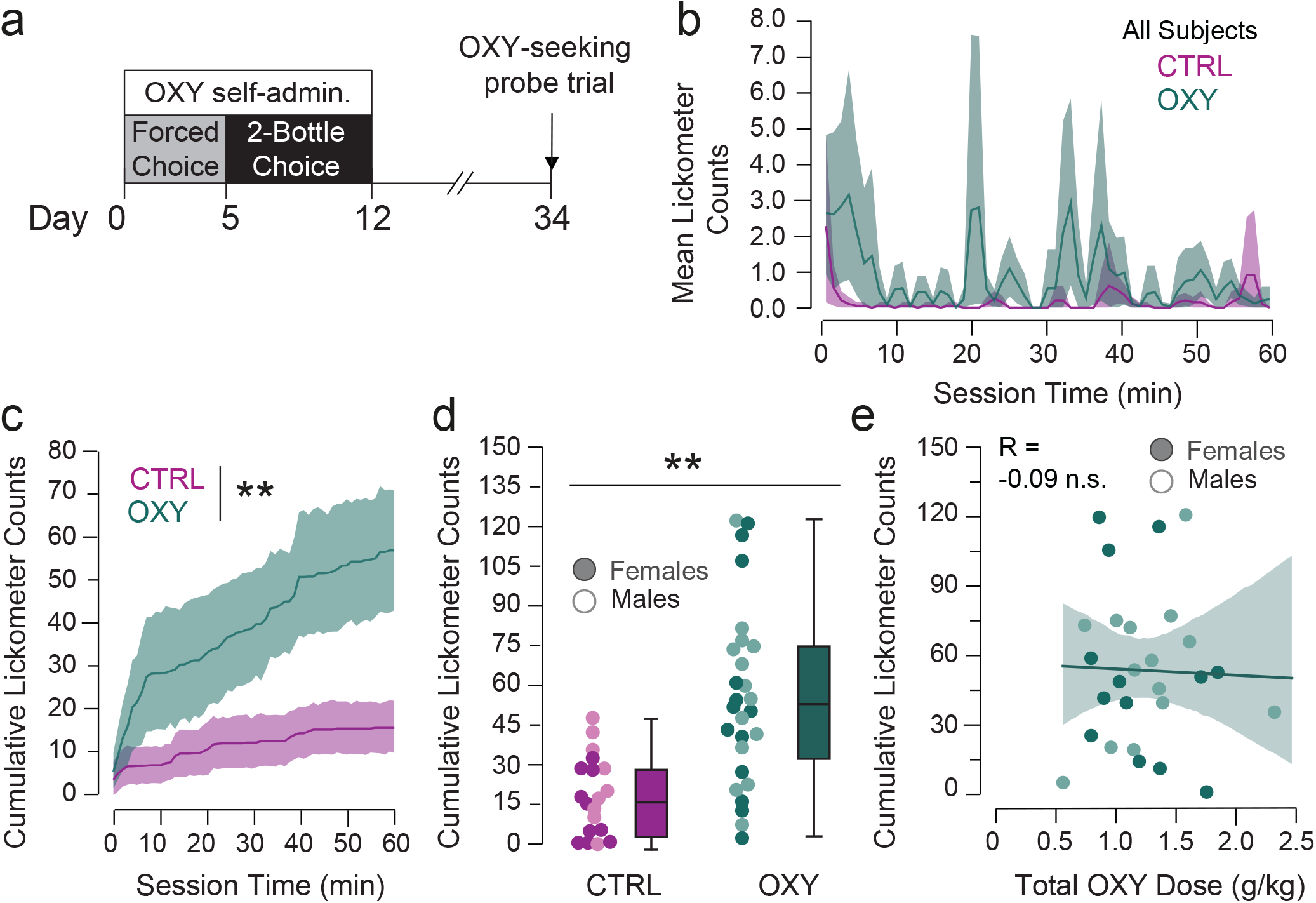
OXY self-administration supports persistent drug seeking behavior in extinction. **(a)** Experimental timeline. **(b-c)** Rolling average time course (a) and cumulative counts **(b)** of photobeam lickometer counts made on empty bottles over the 60 minute probe trial; mice that had self-administered OXY showed higher lickometer interactions relative to controls throughout the session. **(d)** Both male and female mice that had self-administered OXY exhibited higher sipper counts in extinction (CTRL 9M/10F: 18.32 ± 3.48, OXY 13F/13M: 54.78 ± 6.43, F_drug_ = 18.75 p < 0.0001, F_Sex_ = 0.575 p = 0.452). **(e)** Lickometer counts during the seeking probe trial were not significantly correlated with total OXY consumed (p = 0.869). **p<0.01

### Oral OXY self-administration is associated with plasticity of excitatory synapses onto NAcSh Medium Spiny Neurons (MSNs)

Extensive prior work has established that excitatory drive onto MSNs in the nucleus accumbens shell (NAcSh) is potentiated following withdrawal from self-administration of opioids [47, 71-74] and psychostimulants [14, 75, 76]. Critically, this potentiation has been causally linked to persistent cue- and context-induced drug seeking behavior [14, 48, 75, 77-80]. As this potentiation is mediated by the insertion of AMPA receptors into the post-synaptic membrane of accumbal MSNs, we performed whole cell recordings of accumbal MSNs in the NAcSh (Fig 5A,B, Supp Fig 7A,B) to determine whether an analogous form of plasticity would be observed following oral OXY self-administration. Indeed, the AMPA-to-NMDA ratio of electrically-evoked synaptic currents was significantly higher in mice who underwent OXY self-administration and withdrawal relative to control mice, suggesting post-synaptic insertion of AMPA receptors (Fig 5C). Consistent with this interpretation, there were no differences in PPR of electrically-evoked currents between OXY self-administering mice and control mice (Fig 5D), which suggests there was no change in pre-synaptic release onto accumbal MSNs. There was no correlation between total OXY intake and either electrophysiological parameter (Supp Fig 7 C,D). This finding is consistent with prior electrophysiological assays of NAc plasticity following exposure to addictive drugs; potentiation of excitatory transmission onto NAcSh MSNs is preferentially but not selectively expressed onto D1-MSNs [81-83] (but see[75]), while morphine self-administration potentiates excitatory drive onto both D1- and D2-MSNs[47, 73]. Since we used electrical stimulation in wild-type mice, we cannot determine whether these adaptations are specific to a particular excitatory accumbal input, nor whether they occur in both D1- and D2-MSNs within the accumbens, although a bimodal distribution of the AMPA/NMDA ratios raises the possibility that adaptations may be cell-type specific which may be investigated in future studies. Together, these experiments demonstrate that oral OXY self-administration induces behavioral adaptations and associated excitatory synaptic plasticity in the NAcSh that persists into withdrawal.

**Figure 5.**
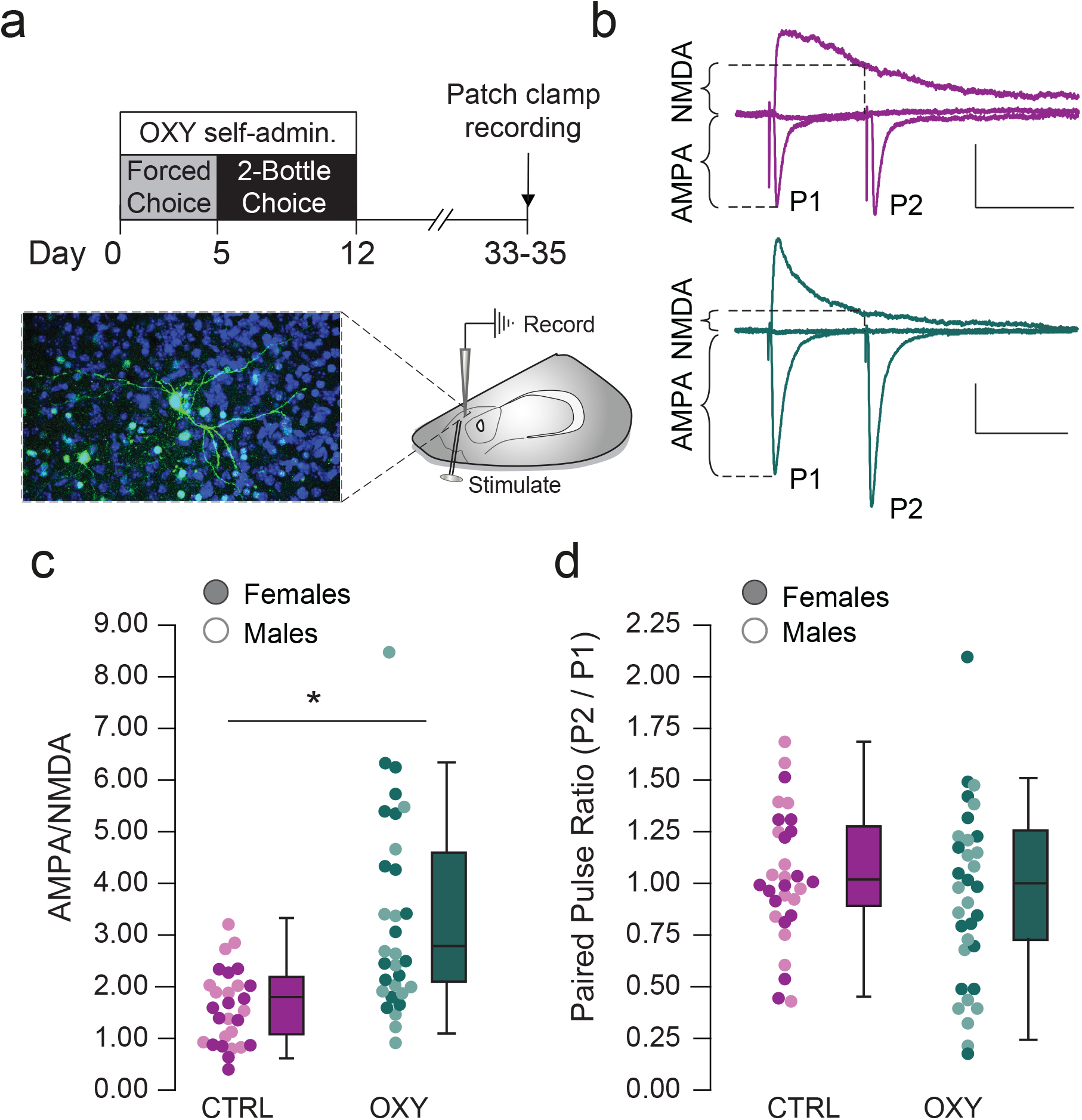
Oral OXY self-administration is associated with plasticity of excitatory synapses onto NAcSh MSNs. **(a)** Experimental timeline and representative alexofluor-488 filled MSN (green), DAPI nuclear stain shown in blue. **(b)** Representative traces showing calculation of AMPA/NMDA and paired pulse ratio in control (top) and OXY withdrawn (bottom) groups; scale bar = 50 pA, 50 ms. **(c)** There was a significant increase in A/N following OXY withdrawal (CTRL female = 1.69 ± 0.19, male = 1.90 ± 0.20, OXY female = 3.56 ± 0.45, male = 3.36 ± 0.53; F_drug_ = 28.62 p < 0.0001). **(d)** PPR was not different between groups (CTRL female = 1.10 ± 0.09, male = 1.01 ± 0.07, OXY female = 0.94 ± 0.12, male = 0.96 ± 0.08; F_drug_ = 1.09, p=0.302). *p<0.05.

### The two-bottle choice paradigm induces aversion-resistant drug consumption in a subset of mice

Another DSM V criteria of substance use disorder is continued use of the substance despite negative consequences. Negative consequences are frequently modeled by a physical punishment such as foot shock [81, 84, 85] or adulteration with aversive tastants, such as quinine[86, 87]. Given the analgesic properties of OXY that could confound responses to foot shock punishment, we introduced increasing concentrations of quinine to the OXY-containing drinking tube, such that the quinine concentration increased by 125 µM every 48 hours (Fig 6A). Both male and female CTRL mice avoided the quinine-containing bottle at the lowest quinine concentration (Fig 6B). By contrast, a subset of OXY mice exhibited quinine-resistant OXY intake, as they persisted in drinking OXY beyond intermediate concentrations of quinine (n=4F/3M out of 15 total mice; Fig 6C, D). There were no sex differences in the AUC of the quinine preference curve (Fig 6E).

**Figure 6.**
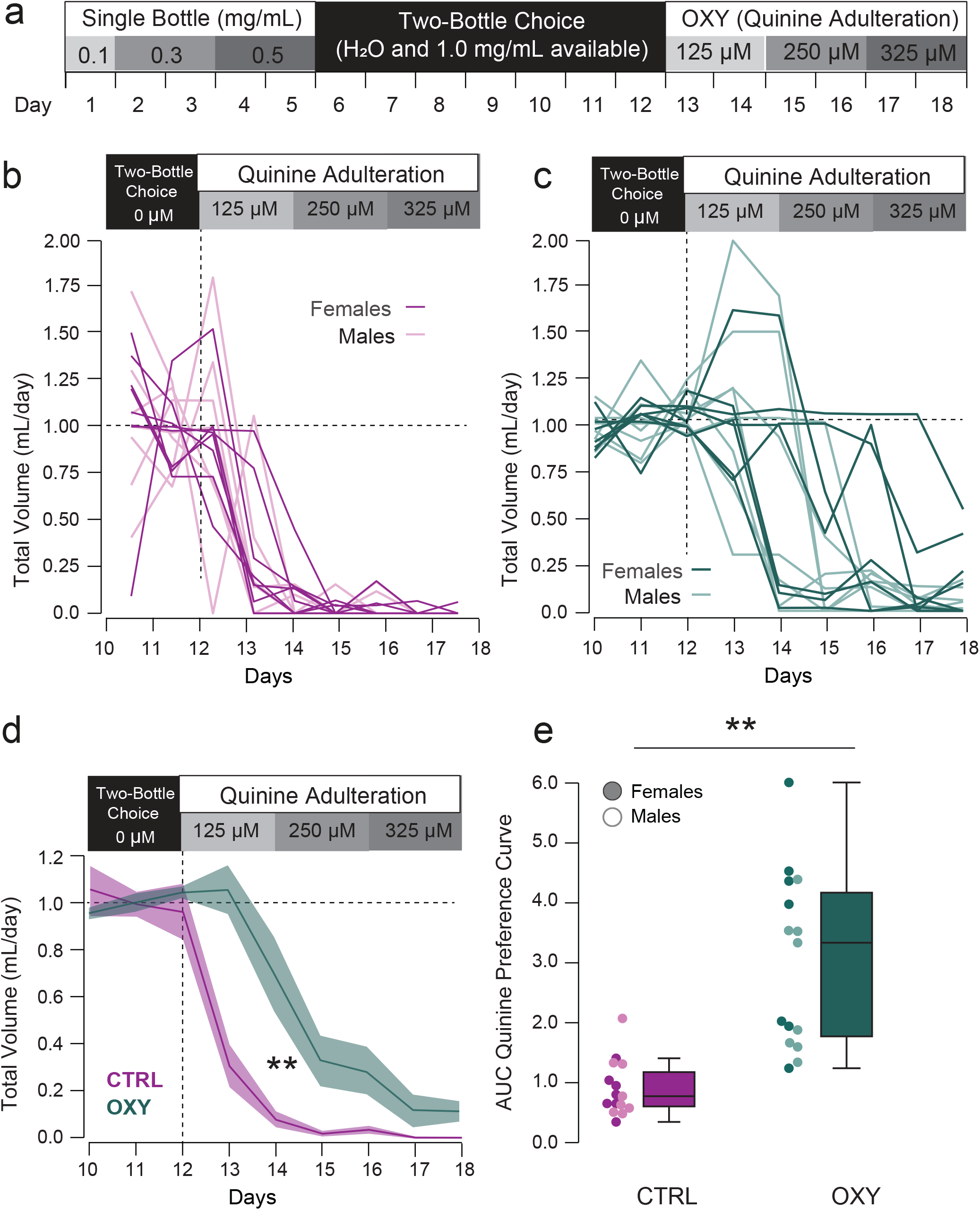
Oral OXY self-administration is resistant to quinine adulteration. **(a)** Experimental schematic. After seven days of two-bottle choice, the OXY-containing bottle (or one H_2_O bottle in CTRL mice) was adulterated with increasing concentrations of quinine. CTRL = 7M/8F, OXY n=8M/7F. **(b-c)** Normalized volume consumed from the quinine-adulterated bottle over the final three days of two bottle choice (before quinine adulteration) and for 6 days of quinine adulteration. **(d-e)** Area under the curve of adulterated quinine drinking volumes. There was a significant effect of Drug (F_drug_ = 27.908, p<0.001), of day (F_day_ = 66.767, p<0.001) and interaction (F_drug•day_ = 7.485, p<0.001), with both male and female OXY mice exhibiting significantly higher AUC relative to CTRL mice (F_sex_ = 0.6896, p = 0.414, CTRL female = 0.862 ± 0.130, male = 0.989 ± 0.201, OXY female = 3.495 ± 0.662, male = 2.704 ± 0.415). **p<0.01

## Discussion

Despite increased awareness of the abuse potential of prescription opioid use and altered prescribing practices, there were still over 142 million prescriptions written for prescription opioids in 2020[88] with opioid-related deaths reaching their highest rate ever in 2021[89]. To develop effective prevention and treatment strategies to treat prescription opioid use disorder, preclinical models that recapitulate behavioral phenomena specific to prescription opioid use are needed. Environmental factors, route of administration, and pattern of intake strongly influence many aspects of drug misuse, which emphasizes the need for preclinical models that recapitulate these features.

One interesting feature of the oral OXY self-administration paradigm characterized here was that considerable individual variability in OXY preference and intake among mice. Similar variability in these factors occurs in people who take prescription opiates. However, at the population level mice escalate their OXY consumption (Fig 1, S1), and alter their pattern of intake (Fig 2, S2, S3) which are two DSM-defined criteria of OUD [11,96,97]. Female mice consumed more OXY (mg/day) during the single bottle phase and higher total dose of OXY (mg/kg) compared to males (Fig 1), consistent with previous observations that female rodents self-administer higher doses of opioids than males [2, 90-94]. While pharmacokinetic influences on sex differences are beyond the scope of our study, plasma concentrations of OXY and its active metabolite oxymorphone, was influenced by sex, presence of sex organs, and feeding conditions, and was exacerbated following repeated administrations (31489746). This likely arises because OXY metabolism is mediated largely by CYP3A4 and CYP2D6 [95, 96], whose activity is modulated by sex hormones and food intake [97, 98]. However, sex effects on OXY pharmacokinetics were either diminished or not observed with intravenous administration [99, 100], highlighting the importance of route of administration for studying sex differences in preclinical studies.

A hallmark DSMV-defined criteria of substance use disorder is the continued use of an addictive substance despite negative consequences. We modeled this feature by adulterating OXY with increasing doses of quinine, which is commonly used to assess compulsive alcohol drinking [101, 102]. We observed high variability in quinine-resistant OXY consumption, with only a subset of mice persisting at the highest quinine doses (Fig 6). This is reminiscent of intravenous opioid or psychostimulant self-administration, where typically only a subpopulation (15-30%) of rodents persist in drug self-administration despite punishment [81, 103-105]. This is critical, as many patients also use opioids as prescribed, with only a subpopulation transitioning to compulsive drug use [104, 106]. This underscores the utility of this model for studying individual differences in aversion-resistant OXY consumption and targeted interventions to modulate this aversion-resistance.

A key challenge in treating patients with OUD is relapse following abstinence. The factors driving relapse in opioid-self-administering patients are complex and are thought to involve a combination of persistent drug craving and avoiding the aversive state of OXY withdrawal [107, 108]. Our results suggest that multiple features and neurobiological substrates relevant to relapse are present with this oral self-administration paradigm. Specifically, drug craving [40] is frequently modeled with reinstatement paradigms, in which an instrumental response previously leading to the addictive drug is assayed under extinction conditions [20, 25, 109, 110]. Here, we experimentally imposed abstinence for three weeks following OXY self-administration [69, 111], to emulate withdrawal imposed by admission to in-patient treatment facilities, or loss of access to prescription or recreational sources [102]. Lickometer interactions which had resulted in the delivery of OXY during the self-administration phase were significantly higher in OXY-self-administering mice relative to controls following protracted withdrawal (Fig 3), consistent with cue-induced OXY-seeking behavior [112-114]. Moreover, OXY self-administering mice exhibited dose-dependent physical withdrawal signs, which persisted (albeit to a lesser extent) into this protracted withdrawal period (Fig 2). In parallel, we found that OXY self-administration resulted in post-synaptically-mediated potentiation of excitatory transmission into NAc MSNs (Fig 5), which has been causally linked to both the persistence of reinstatement behavior and physical symptoms following withdrawal from addictive drugs [14, 48, 80, 115-119]. It is important to note that our electrophysiology studies used electrical stimulation to induce excitatory synaptic currents and were performed in wild-type mice, and thus we do not know which specific accumbal populations or excitatory inputs are potentiated following oral OXY self-administration, although this represents an important future application of this preclinical model. Taken together, these findings underscore the utility of this paradigm for studying diverse factors contributing to opioid relapse (i.e. emergence of withdrawal symptoms and craving), along with neural adaptations that may underlie the persistence of these behavioral symptoms.

The high-throughput nature of the model characterized here enables high-powered studies that facilitate understanding of individual variability in drug taking behavior, reinstatement and persistence of drug taking despite negative consequences, while conserving the route of administration, environmental factors pattern of access specifically relevant to prescription opioid misuse. Future work is needed to understand the neural substrates underlying the emergence of prescription opioid reinstatement or aversion-resistant intake, or to inform strategies to manage prescription opioid abuse, including optimizing tapering schedules or assaying pharmacological adjuvants to manage withdrawal symptoms. OXY is primarily prescribed for the treatment of pain and is frequently used in combination with other prescription medications or recreationally used substances. To this end, this model could be used in future studies investigating how pre-existing chronic or post-surgical pain, or polysubstance use modulates patterns of OXY-intake, and the consequences of this intake for the expression of addiction-relevant behaviors. The current results present a first characterization of a preclinical model that could be applied towards these goals.

## Acknowledgements

We thank M.C. Stander and E. Godynyuk for excellent technical help. We also J.J. Choong, S. Golden and M. Mathis for their commitment to open-source tools and for support with supervised behavioral classification (SimBA) and markerless pose estimation (DeepLabCut™), respectively. We thank MCCI Corporation (Ithaca NY) and Terrence Moore for development of wireless in-cage activity and environmental sensors and the IoT cloud solution.

## Author Contributions

RAS, YHC, RP, EC, JT, YMV, DKW and JTS performed behavioral experiments. JL, JRT, BC, KA and MCC performed patch-clamp experiments. MCC, AVK developed home cage-based lickometer devices for self-administration, AVK developed PIR sensors and cloud-based device monitoring, TE developed custom software for analysis. RAS, BC, and MCC analyzed data. RG, BC, AVK, and MCC supervised work. MCC, RAS wrote the manuscript with input from all authors.

## References

1. Vowles, K.E., et al., Rates of opioid misuse, abuse, and addiction in chronic pain: a systematic review and data synthesis. Pain, 2015. 156(4): p. 569–576.

2. Phillips, A.G., et al., Oral prescription opioid-seeking behavior in male and female mice. Addict Biol, 2020. 25(6): p. e12828.

3. Enga, R.M., et al., Oxycodone physical dependence and its oral self-administration in C57BL/6J mice. Eur J Pharmacol, 2016. 789: p. 75–80.

4. Fulenwider, H.D., et al., Sex differences in oral oxycodone self-administration and stress-primed reinstatement in rats. Addict Biol, 2020. 25(6): p. e12822.

5. Kirsh, K., J. Peppin, and J. Coleman, Characterization of Prescription Opioid Abuse in the United States: Focus on Route of Administration. Journal of Pain & Palliative Care Pharmacotherapy, 2012. 26(4): p. 348–361.

6. Leow, K.P., et al., Comparative Oxycodone Pharmacokinetics in Humans After Intravenous, Oral, and Rectal Administration. Therapeutic Drug Monitoring, 1992. 14(6): p. 479–484.

7. Poyhia, R., et al., The pharmacokinetics and metabolism of oxycodone after intramuscular and oral administration to healthy subjects. Br J Clin Pharmacol, 1992. 33(6): p. 617–21.

8. Lugo, R.A. and S.E. Kern, The pharmacokinetics of oxycodone. J Pain Palliat Care Pharmacother, 2004. 18(4): p. 17–30.

9. Streisand, J.B., et al., Absorption and bioavailability of oral transmucosal fentanyl citrate. Anesthesiology, 1991. 75(2): p. 223–9.

10. Drewes, A.M., et al., Differences between opioids: pharmacological, experimental, clinical and economical perspectives. Br J Clin Pharmacol, 2013. 75(1): p. 60–78.

11. Goodman, L.S., Goodman and Gilman’s the pharmacological basis of therapeutics. Vol. 1549. 1996: McGraw-Hill New York.

12. Yoburn, B.C., et al., Supersensitivity to opioid analgesics following chronic opioid antagonist treatment: relationship to receptor selectivity. Pharmacol Biochem Behav, 1995. 51(2-3): p. 535–9.

13. Smith, H.S., Opioid metabolism. Mayo Clin Proc, 2009. 84(7): p. 613–24.

14. Conrad, K.L., et al., Formation of accumbens GluR2-lacking AMPA receptors mediates incubation of cocaine craving. Nature, 2008. 454(7200): p. 118–121.

15. Volkow, N.D., et al., Addiction: decreased reward sensitivity and increased expectation sensitivity conspire to overwhelm the brain’s control circuit. Bioessays, 2010. 32(9): p. 748–55.

16. Balster, R.L. and C.R. Schuster, Fixed-interval schedule of cocaine reinforcement: effect of dose and infusion duration. J Exp Anal Behav, 1973. 20(1): p. 119–29.

17. Jacob, J.C., et al., Ethanol Reversal of Tolerance to the Antinociceptive Effects of Oxycodone and Hydrocodone. Journal of Pharmacology and Experimental Therapeutics, 2017. 362(1): p. 45–52.

18. Macenski, M.J. and R.A. Meisch, Oral Drug Reinforcement Studies With Laboratory Animals: Applications and Implications for Understanding Drug-Reinforced Behavior. Current Directions in Psychological Science, 1994. 3(1): p. 22–27.

19. Venniro, M., et al., Improving translation of animal models of addiction and relapse by reverse translation. Nature Reviews Neuroscience, 2020. 21(11): p. 625–643.

20. Venniro, M., D. Caprioli, and Y. Shaham, Animal models of drug relapse and craving: From drug priming-induced reinstatement to incubation of craving after voluntary abstinence. Prog Brain Res, 2016. 224: p. 25–52.

21. Panlilio, L.V., S.J. Weiss, and C.W. Schindler, Effects of compounding drug-related stimuli: escalation of heroin self-administration. Journal of the experimental analysis of behavior, 2000. 73(2): p. 211–224.

22. Perry, C.J., et al., Role of cues and contexts on drug-seeking behaviour. British journal of pharmacology, 2014. 171(20): p. 4636–4672.

23. Wakabayashi, K.T., et al., Rats markedly escalate their intake and show a persistent susceptibility to reinstatement only when cocaine is injected rapidly. The Journal of neuroscience : the official journal of the Society for Neuroscience, 2010. 30(34): p. 11346–11355.

24. Meisch, R.A. and M.E. Carroll, Oral Drug Self-Administration: Drugs as Reinforcers, in Methods of Assessing the Reinforcing Properties of Abused Drugs. 1987, Springer New York. p. 143–160.

25. Reiner, D.J., et al., Relapse to opioid seeking in rat models: behavior, pharmacology and circuits. Neuropsychopharmacology : official publication of the American College of Neuropsychopharmacology, 2019. 44(3): p. 465–477.

26. Calipari, E.S., et al., Intermittent cocaine self-administration produces sensitization of stimulant effects at the dopamine transporter. J Pharmacol Exp Ther, 2014. 349(2): p. 192–8.

27. Calipari, E.S., et al., Temporal pattern of cocaine intake determines tolerance vs sensitization of cocaine effects at the dopamine transporter. Neuropsychopharmacology, 2013. 38(12): p. 2385–92.

28. Lefevre, E.M., et al., Interruption of continuous opioid exposure exacerbates drug-evoked adaptations in the mesolimbic dopamine system. Neuropsychopharmacology : official publication of the American College of Neuropsychopharmacology, 2020. 45(11): p. 1781–1792.

29. Cahill, C.M., Opioid dose regimen shapes mesolimbic adaptations. Neuropsychopharmacology : official publication of the American College of Neuropsychopharmacology, 2020. 45(11): p. 1777–1778.

30. Zimmer, B.A., E.B. Oleson, and D.C. Roberts, The motivation to self-administer is increased after a history of spiking brain levels of cocaine. Neuropsychopharmacology, 2012. 37(8): p. 1901–10.

31. Caprioli, D., et al., Modeling the role of environment in addiction. Progress in Neuro-Psychopharmacology and Biological Psychiatry, 2007. 31(8): p. 1639–1653.

32. Badiani, A., et al., Opiate versus psychostimulant addiction: the differences do matter. Nature reviews. Neuroscience, 2011. 12(11): p. 685–700.

33. Badiani, A. and P. Spagnolo, Role of Environmental Factors in Cocaine Addiction. Current Pharmaceutical Design, 2013. 19(40): p. 6996–7008.

34. Caprioli, D., et al., Ambience and Drug Choice: Cocaine- and Heroin-Taking as a Function of Environmental Context in Humans and Rats. Biological Psychiatry, 2009. 65(10): p. 893–899.

35. Montanari, C., et al., G.3 - THE ROLE OF ENVIRONMENTAL CONTEXT IN THE VULNERABILITY TO RELAPSE INTO HEROIN AND COCAINE ADDICTION. Behavioural Pharmacology, 2013. 24: p. e56–e57.

36. Drouin, C., et al., Alpha1b-adrenergic receptors control locomotor and rewarding effects of psychostimulants and opiates. J Neurosci, 2002. 22(7): p. 2873–84.

37. Borg, P.J. and D.A. Taylor, Voluntary oral morphine self-administration in rats: effect of haloperidol or ondansetron. Pharmacol Biochem Behav, 1994. 47(3): p. 633–46.

38. Grim, T.W., et al., The effect of quinine in two bottle choice procedures in C57BL6 mice: Opioid preference, somatic withdrawal, and pharmacokinetic outcomes. Drug Alcohol Depend, 2018. 191: p. 195–202.

39. Winterdahl, M., et al., Sucrose intake lowers μ-opioid and dopamine D2/3 receptor availability in porcine brain. Scientific reports, 2019. 9(1): p. 16918–16918.

40. Pickens, C.L., et al., Neurobiology of the incubation of drug craving. Trends in neurosciences, 2011. 34(8): p. 411–420.

41. Bainier, C., et al., Circadian rhythms of hedonic drinking behavior in mice. Neuroscience, 2017. 349: p. 229–238.

42. Hajnal, A., G.P. Smith, and R. Norgren, Oral sucrose stimulation increases accumbens dopamine in the rat. American Journal of Physiology-Regulatory, Integrative and Comparative Physiology, 2004. 286(1): p. R31–R37.

43. Hoebel, B.G., et al., Natural addiction: a behavioral and circuit model based on sugar addiction in rats. Journal of addiction medicine, 2009. 3(1): p. 33–41.

44. Rada, P., N.M. Avena, and B.G. Hoebel, Daily bingeing on sugar repeatedly releases dopamine in the accumbens shell. Neuroscience, 2005. 134(3): p. 737–744.

45. Spangler, R., et al., Opiate-like effects of sugar on gene expression in reward areas of the rat brain. Molecular Brain Research, 2004. 124(2): p. 134–142.

46. Diagnostic and Statistical Manual of Mental Disorders, 5th Edition. 2013, American Psychiatric Publishing, Inc.

47. Madayag, A.C., et al., Cell-type and region-specific nucleus accumbens AMPAR plasticity associated with morphine reward, reinstatement, and spontaneous withdrawal. Brain structure & function, 2019. 224(7): p. 2311–2324.

48. Mameli, M., et al., Cocaine-evoked synaptic plasticity: persistence in the VTA triggers adaptations in the NAc. Nature Neuroscience, 2009. 12(8): p. 1036–1041.

49. Creed, M.C. and C. Lüscher, Drug-evoked synaptic plasticity: beyond metaplasticity. Current Opinion in Neurobiology, 2013. 23(4): p. 553–558.

50. Matikainen-Ankney, B.A., et al., Rodent Activity Detector (RAD), an Open Source Device for Measuring Activity in Rodent Home Cages. eNeuro, 2019. 6(4): p. ENEURO.0160-19.2019.

51. Godynyuk, E., et al., An Open-Source, Automated Home-Cage Sipper Device for Monitoring Liquid Ingestive Behavior in Rodents. eNeuro, 2019. 6(5): p. ENEURO.0292-19.2019.

52. Nilsson, S.R.O., et al., Simple Behavioral Analysis (SimBA) – an open source toolkit for computer classification of complex social behaviors in experimental animals. 2020, Cold Spring Harbor Laboratory.

53. Abraham, A., et al., Machine learning for neuroimaging with scikit-learn. Frontiers in neuroinformatics, 2014. 8: p. 14–14.

54. Zelinsky, A., Learning OpenCV---Computer Vision with the OpenCV Library (Bradski,G.R. et al.; 2008)[On the Shelf]. IEEE Robotics & Automation Magazine, 2009. 16(3): p. 100–100.

55. new ffmpeg, in SciVee. 2009, SciVee, Inc.

56. Lemaitre, G., F. Nogueira, and C.K. Aridas, Imbalanced-learn: A Python Toolbox to Tackle the Curse of Imbalanced Datasets in Machine Learning. Journal of Machine Learning Research, 2017. 18.

57. Liu, Y.-l., et al., Effects of l-tetrahydropalmatine on locomotor sensitization to oxycodone in mice. Acta Pharmacologica Sinica, 2005. 26(5): p. 533–538.

58. Niikura, K., et al., Oxycodone-induced conditioned place preference and sensitization of locomotor activity in adolescent and adult mice. Pharmacology, biochemistry, and behavior, 2013. 110: p. 112–116.

59. Bryant, C.D., et al., The heritability of oxycodone reward and concomitant phenotypes in a LG/J × SM/J mouse advanced intercross line. Addiction biology, 2014. 19(4): p. 552–561.

60. Gutierrez, A., et al., Female rats self-administer heroin by vapor inhalation. Pharmacology, biochemistry, and behavior, 2020. 199: p. 173061–173061.

61. Sabeti, J., G.A. Gerhardt, and N.R. Zahniser, Individual Differences in Cocaine-Induced Locomotor Sensitization in Low and High Cocaine Locomotor-Responding Rats Are Associated with Differential Inhibition of Dopamine Clearance in Nucleus Accumbens. Journal of Pharmacology and Experimental Therapeutics, 2003. 305(1): p. 180–190.

62. Fennell, A.M., et al., Phasic Dopamine Release Magnitude Tracks Individual Differences in Sensitization of Locomotor Response following a History of Nicotine Exposure. Scientific reports, 2020. 10(1): p. 173–173.

63. De Vries, T.J., A.R. Cools, and T.S. Shippenberg, Infusion of a D-1 receptor agonist into the nucleus accumbens enhances cocaine-induced behavioural sensitization. NeuroReport, 1998. 9(8): p. 1763–1768.

64. Reeves, K.C., et al., Mu opioid receptors on vGluT2-expressing glutamatergic neurons modulate opioid reward. Addiction biology, 2021. 26(3): p. e12942–e12942.

65. Collins, D., et al., Sex differences in responsiveness to the prescription opioid oxycodone in mice. Pharmacology Biochemistry and Behavior, 2016. 148: p. 99–105.

66. Grim, T.W., et al., A G protein signaling-biased agonist at the mu-opioid receptor reverses morphine tolerance while preventing morphine withdrawal. Neuropsychopharmacology, 2020. 45(2): p. 416–425.

67. Schulteis, G., et al., Relative sensitivity to naloxone of multiple indices of opiate withdrawal: a quantitative dose-response analysis. J Pharmacol Exp Ther, 1994. 271(3): p. 1391–8.

68. Schreiber, S., et al., Mianserin and trazodone significantly attenuate the intensity of opioid withdrawal symptoms in mice. Addict Biol, 2003. 8(1): p. 107–14.

69. Wolf, M.E., Synaptic mechanisms underlying persistent cocaine craving. Nature reviews. Neuroscience, 2016. 17(6): p. 351–365.

70. Bossert, J.M., et al., The reinstatement model of drug relapse: recent neurobiological findings, emerging research topics, and translational research. Psychopharmacology, 2013. 229(3): p. 453–476.

71. Graziane, N.M., et al., Opposing mechanisms mediate morphine- and cocaine-induced generation of silent synapses. Nature neuroscience, 2016. 19(7): p. 915–925.

72. Russell, S.E., et al., Nucleus Accumbens AMPA Receptors Are Necessary for Morphine-Withdrawal-Induced Negative-Affective States in Rats. The Journal of neuroscience : the official journal of the Society for Neuroscience, 2016. 36(21): p. 5748–5762.

73. Zhu, Y., et al., A thalamic input to the nucleus accumbens mediates opiate dependence. Nature, 2016. 530(7589): p. 219–222.

74. Hearing, M.C., et al., Reversal of morphine-induced cell-type-specific synaptic plasticity in the nucleus accumbens shell blocks reinstatement. Proceedings of the National Academy of Sciences of the United States of America, 2016. 113(3): p. 757–762.

75. Terrier, J., C. Lüscher, and V. Pascoli, Cell-Type Specific Insertion of GluA2-Lacking AMPARs with Cocaine Exposure Leading to Sensitization, Cue-Induced Seeking, and Incubation of Craving. Neuropsychopharmacology : official publication of the American College of Neuropsychopharmacology, 2016. 41(7): p. 1779–1789.

76. Murray, C.H., et al., AMPA receptor and metabotropic glutamate receptor 1 adaptations in the nucleus accumbens core during incubation of methamphetamine craving. Neuropsychopharmacology : official publication of the American College of Neuropsychopharmacology, 2019. 44(9): p. 1534–1541.

77. Hollander, J.A. and R.M. Carelli, Abstinence from Cocaine Self-Administration Heightens Neural Encoding of Goal-Directed Behaviors in the Accumbens. Neuropsychopharmacology, 2005. 30(8): p. 1464–1474.

78. Pascoli, V., et al., Contrasting forms of cocaine-evoked plasticity control components of relapse. Nature, 2014. 509(7501): p. 459–464.

79. Ma, Y.-Y., et al., Bidirectional modulation of incubation of cocaine craving by silent synapse-based remodeling of prefrontal cortex to accumbens projections. Neuron, 2014. 83(6): p. 1453–1467.

80. Ortinski, P.I., et al., Temporally dependent changes in cocaine-induced synaptic plasticity in the nucleus accumbens shell are reversed by D1-like dopamine receptor stimulation. Neuropsychopharmacology : official publication of the American College of Neuropsychopharmacology, 2012. 37(7): p. 1671–1682.

81. Pascoli, V., et al., Sufficiency of Mesolimbic Dopamine Neuron Stimulation for the Progression to Addiction. Neuron, 2015. 88(5): p. 1054–1066.

82. Creed, M., V.J. Pascoli, and C. Lüscher, Refining deep brain stimulation to emulate optogenetic treatment of synaptic pathology. Science, 2015. 347(6222): p. 659–664.

83. Pascoli, V., M. Turiault, and C. Lüscher, Reversal of cocaine-evoked synaptic potentiation resets drug-induced adaptive behaviour. Nature, 2011. 481(7379): p. 71–75.

84. Pelloux, Y., B.J. Everitt, and A. Dickinson, Compulsive drug seeking by rats under punishment: effects of drug taking history. Psychopharmacology, 2007. 194(1): p. 127–137.

85. Farrell, M.R., et al., Ventral pallidum is essential for cocaine relapse after voluntary abstinence in rats. Neuropsychopharmacology : official publication of the American College of Neuropsychopharmacology, 2019. 44(13): p. 2174–2185.

86. Hopf, F.W. and H.M.B. Lesscher, Rodent models for compulsive alcohol intake. Alcohol (Fayetteville, N.Y.), 2014. 48(3): p. 253–264.

87. McCane, A.M., et al., Differential effects of quinine adulteration of alcohol on seeking and drinking. Alcohol (Fayetteville, N.Y.), 2021. 92: p. 73–80.

88. Prevention, C.f.D.C.a. U.S. Opioid Dispensing Rate Maps. 2021 [cited 2022 2022]; Available from: https://www.cdc.gov/drugoverdose/rxrate-maps/index.html.

89. Association, A.M. Issue brief: Nation’s drug-related overdose and death epidemic continues to worsen. 2022; Available from: https://www.ama-assn.org/system/files/issue-brief-increases-in-opioid-related-overdose.pdf.

90. Zanni, G., et al., Female and male rats readily consume and prefer oxycodone to water in a chronic, continuous access, two-bottle oral voluntary paradigm. Neuropharmacology, 2020. 167: p. 107978.

91. Alexander, B.K., R.B. Coambs, and P.F. Hadaway, The effect of housing and gender on morphine self-administration in rats. Psychopharmacology, 1978. 58(2): p. 175–179.

92. Hadaway, P.F., et al., The effect of housing and gender on preference for morphine-sucrose solutions in rats. Psychopharmacology, 1979. 66(1): p. 87–91.

93. Klein, L.C., E.J. Popke, and N.E. Grunberg, Sex differences in effects of predictable and unpredictable footshock on fentanyl self-administration in rats. Experimental and Clinical Psychopharmacology, 1997. 5(2): p. 99–106.

94. Moussawi, K., et al., Fentanyl vapor self-administration model in mice to study opioid addiction. Science advances, 2020. 6(32): p. eabc0413–eabc0413.

95. Samer, C.F., et al., The effects of CYP2D6 and CYP3A activities on the pharmacokinetics of immediate release oxycodone. British journal of pharmacology, 2010. 160(4): p. 907–918.

96. Grönlund, J., et al., Exposure to oral oxycodone is increased by concomitant inhibition of CYP2D6 and 3A4 pathways, but not by inhibition of CYP2D6 alone. British journal of clinical pharmacology, 2010. 70(1): p. 78–87.

97. Zanger, U.M. and M. Schwab, Cytochrome P450 enzymes in drug metabolism: Regulation of gene expression, enzyme activities, and impact of genetic variation. Pharmacology & Therapeutics, 2013. 138(1): p. 103–141.

98. Soldin, O.P. and D.R. Mattison, Sex differences in pharmacokinetics and pharmacodynamics. Clinical pharmacokinetics, 2009. 48(3): p. 143–157.

99. Chan, S., et al., SEX DIFFERENCES IN THE PHARMACOKINETICS, OXIDATIVE METABOLISM AND ORAL BIOAVAILABILITY OF OXYCODONE IN THE SPRAGUE-DAWLEY RAT. Clinical and Experimental Pharmacology and Physiology, 2008. 35(3): p. 295–302.

100. Mavrikaki, M., et al., Oxycodone self-administration in male and female rats. Psychopharmacology, 2017. 234(6): p. 977–987.

101. Darevsky, D. and F.W. Hopf, Behavioral indicators of succeeding and failing under higher-challenge compulsion-like alcohol drinking in rat. Behavioural brain research, 2020. 393: p. 112768–112768.

102. Fulenwider, H.D., et al., Sex Differences in Aversion-Resistant Ethanol Intake in Mice. Alcohol and Alcoholism, 2019. 54(4): p. 345–352.

103. Kasanetz, F., et al., Transition to Addiction Is Associated with a Persistent Impairment in Synaptic Plasticity. Science, 2010. 328(5986): p. 1709–1712.

104. Pascoli, V., et al., Stochastic synaptic plasticity underlying compulsion in a model of addiction. Nature, 2018. 564(7736): p. 366–371.

105. Deroche-Gamonet, V.r., D. Belin, and P.V. Piazza, Evidence for Addiction-like Behavior in the Rat. Science, 2004. 305(5686): p. 1014–1017.

106. Anthony, J.C., L.A. Warner, and R.C. Kessler, Comparative epidemiology of dependence on tobacco, alcohol, controlled substances, and inhalants: Basic findings from the National Comorbidity Survey. Experimental and Clinical Psychopharmacology, 1994. 2(3): p. 244–268.

107. Shurman, J., G.F. Koob, and H.B. Gutstein, Opioids, pain, the brain, and hyperkatifeia: a framework for the rational use of opioids for pain. Pain medicine (Malden, Mass.), 2010. 11(7): p. 1092–1098.

108. Koob, G.F., Neurobiology of Opioid Addiction: Opponent Process, Hyperkatifeia, and Negative Reinforcement. Biological Psychiatry, 2020. 87(1): p. 44–53.

109. Marchant, N.J., et al., Context-induced relapse after extinction versus punishment: similarities and differences. Psychopharmacology, 2019. 236(1): p. 439–448.

110. Shaham, Y., et al., The reinstatement model of drug relapse: history, methodology and major findings. Psychopharmacology (Berl), 2003. 168(1-2): p. 3–20.

111. Bossert, J.M., et al., Role of mu, but not delta or kappa, opioid receptors in context-induced reinstatement of oxycodone seeking. European Journal of Neuroscience, 2018. 50(3): p. 2075–2085.

112. Neisewander, J.L., et al., Fos protein expression and cocaine-seeking behavior in rats after exposure to a cocaine self-administration environment. J Neurosci, 2000. 20(2): p. 798–805.

113. Grimm, J.W., et al., Neuroadaptation. Incubation of cocaine craving after withdrawal. Nature, 2001. 412(6843): p. 141–2.

114. Wolf, M.E., Synaptic mechanisms underlying persistent cocaine craving. Nat Rev Neurosci, 2016. 17(6): p. 351–65.

115. Scofield, M.D., et al., The Nucleus Accumbens: Mechanisms of Addiction across Drug Classes Reflect the Importance of Glutamate Homeostasis. Pharmacological reviews, 2016. 68(3): p. 816–871.

116. McCutcheon, J.E., et al., Calcium-permeable AMPA receptors are present in nucleus accumbens synapses after prolonged withdrawal from cocaine self-administration but not experimenter-administered cocaine. The Journal of neuroscience : the official journal of the Society for Neuroscience, 2011. 31(15): p. 5737–5743.

117. Jedynak, J., et al., Cocaine and Amphetamine Induce Overlapping but Distinct Patterns of AMPAR Plasticity in Nucleus Accumbens Medium Spiny Neurons. Neuropsychopharmacology : official publication of the American College of Neuropsychopharmacology, 2016. 41(2): p. 464–476.

118. Ebner, S.R., et al., Extinction and Reinstatement of Cocaine-seeking in Self-administering Mice is Associated with Bidirectional AMPAR-mediated Plasticity in the Nucleus Accumbens Shell. Neuroscience, 2018. 384: p. 340–349.

119. Lüscher, C. and R.C. Malenka, Drug-evoked synaptic plasticity in addiction: from molecular changes to circuit remodeling. Neuron, 2011. 69(4): p. 650–663.

