## Supplemental_Figures for "Oral oxycodone self-administration leads to features of opioid addiction in male and female mice"

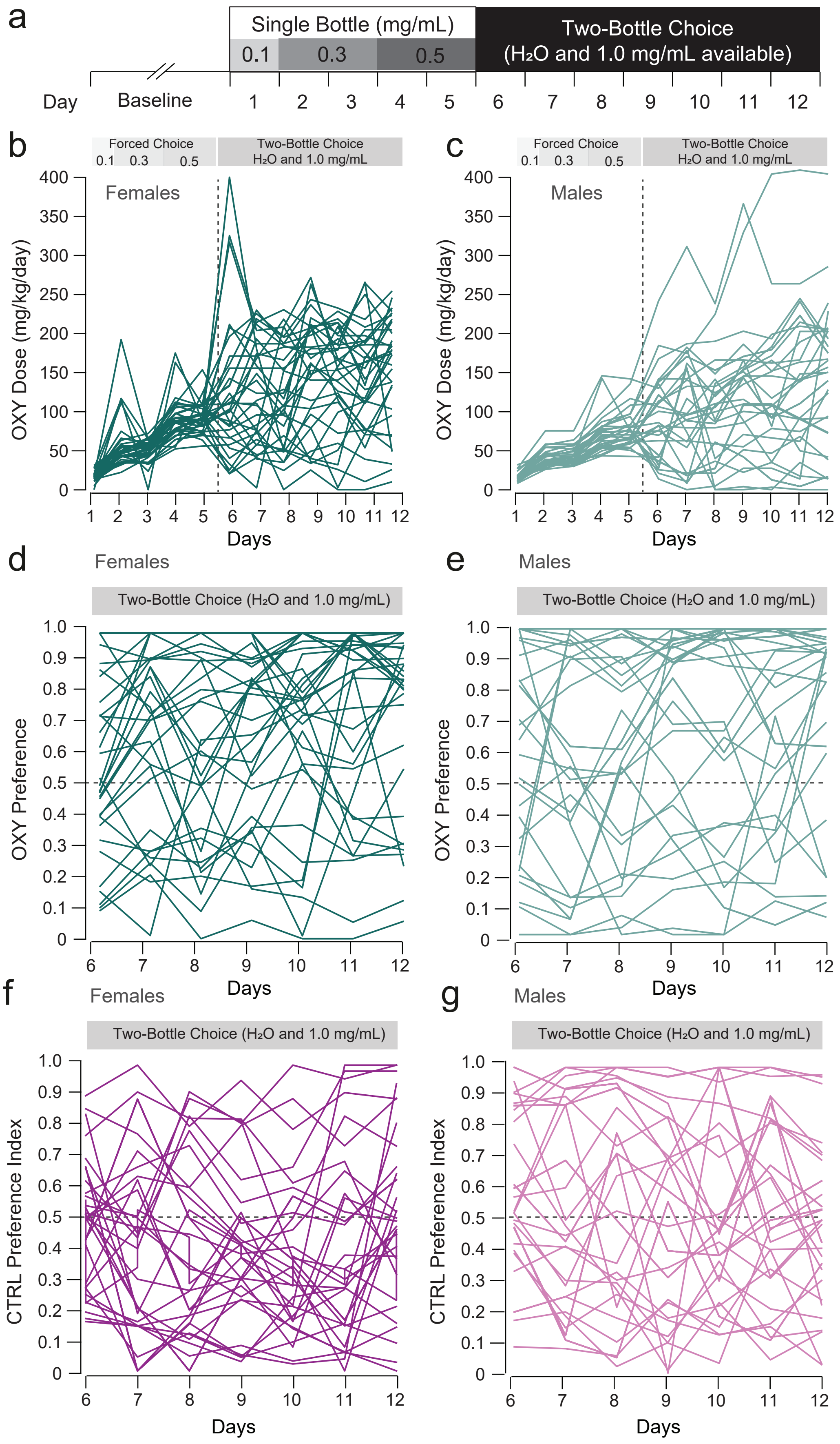


**Supplemental Figure 1. Individual data for oxycodone dose and preference.**

**Supplemental Figure 1. Individual data for OXY dose and preference. a)** Experimental Schematic. **(b-c)** Escalation of OXY dose for individual mice; dose is defined as [*mg OXY consumed / body weight*]. **(d-e)** Preference for OXY for individual mice; preference is defined as [*consumed volume of OXY / total volume of fluid consumption*]. **(f-g)** Preference for the right $H_2O$-containing bottle for individual mice; preference is defined as [*consumed volume from right bottle / total volume of fluid consumption*].

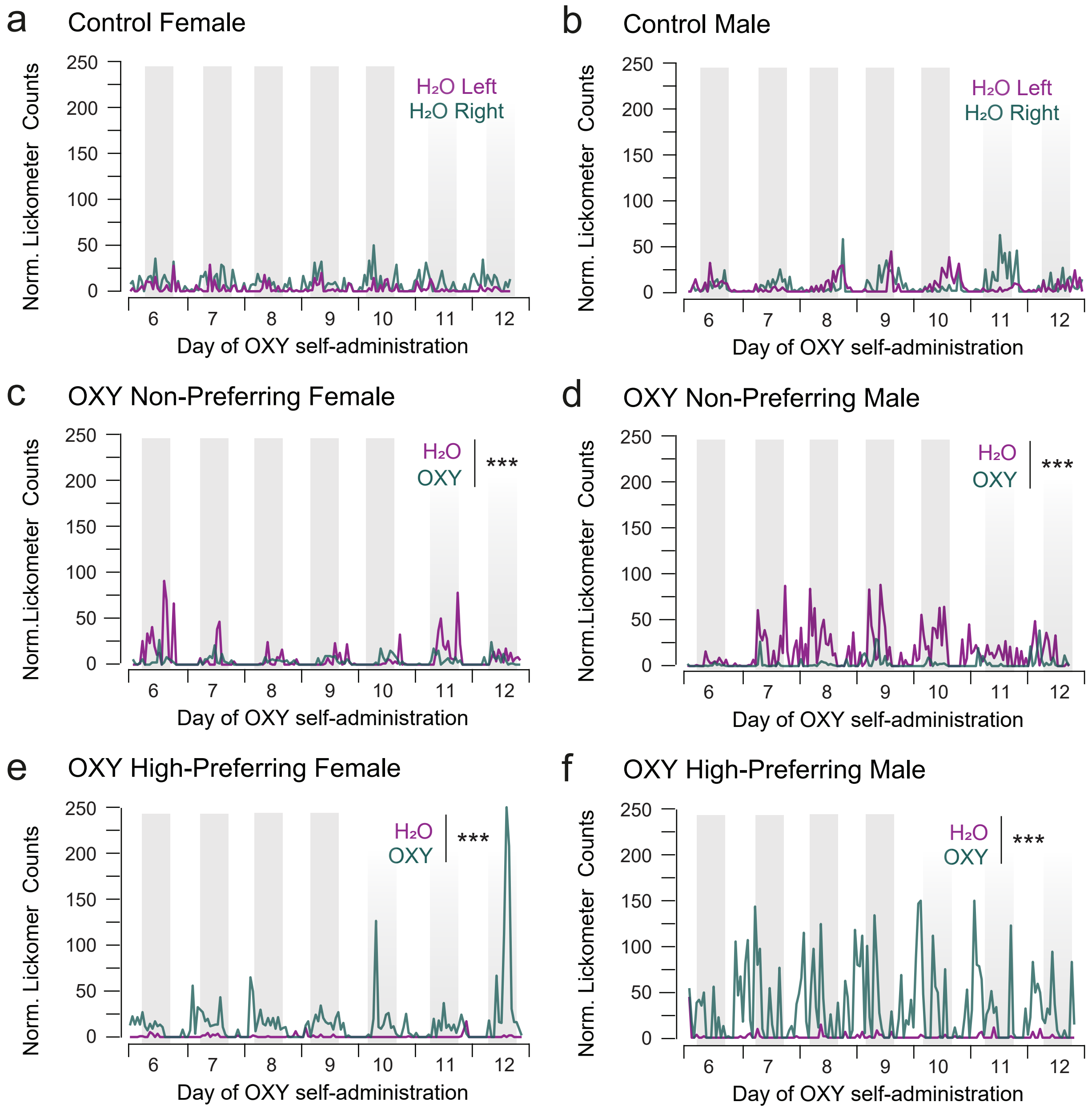


**Supplemental Figure 2. Individual examples of lickometer counts from male and female mice.**

**Supplemental Figure 2. Individual examples of lickometer counts from male and female mice.** Individual lickometer photo-interrupter counts for example control subject are shown. **(a-b)** Counts made on the left and right bottles are shown for controls are shown separately (n = CTRL: 33F/34M, OXY: 35F/33M). Counts made on the OXY-filled and water-filled bottles are shown in both **(c-d)** Non-Preferring (n = 4F/4M) and High-Preferring (n =7F, 6M) and **(e-f)** High-Preferring mice. ***p < 0.001

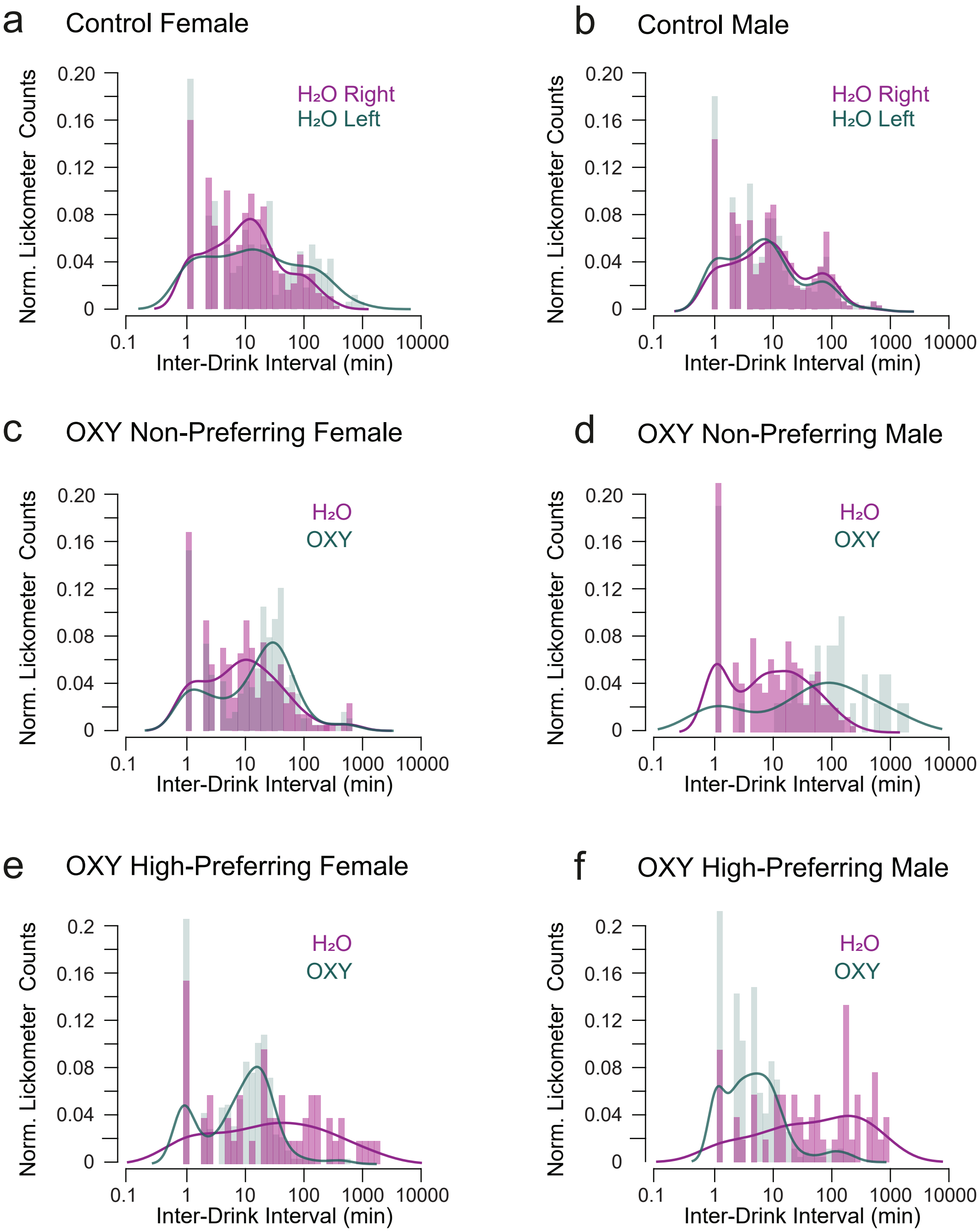


**Supplemental Figure 3. Individual examples of inter-drink-intervals from male and female mice.**

**Supplemental Figure 3. Individual examples of inter-drink-intervals from male and female mice.** Inter-drink intervals from lickometer devices in (a,b) CTRL males and females, (c,d) non-preferring mice (n = 4F/4M), and (e,f) high preferring mice (n =7F, 6M).

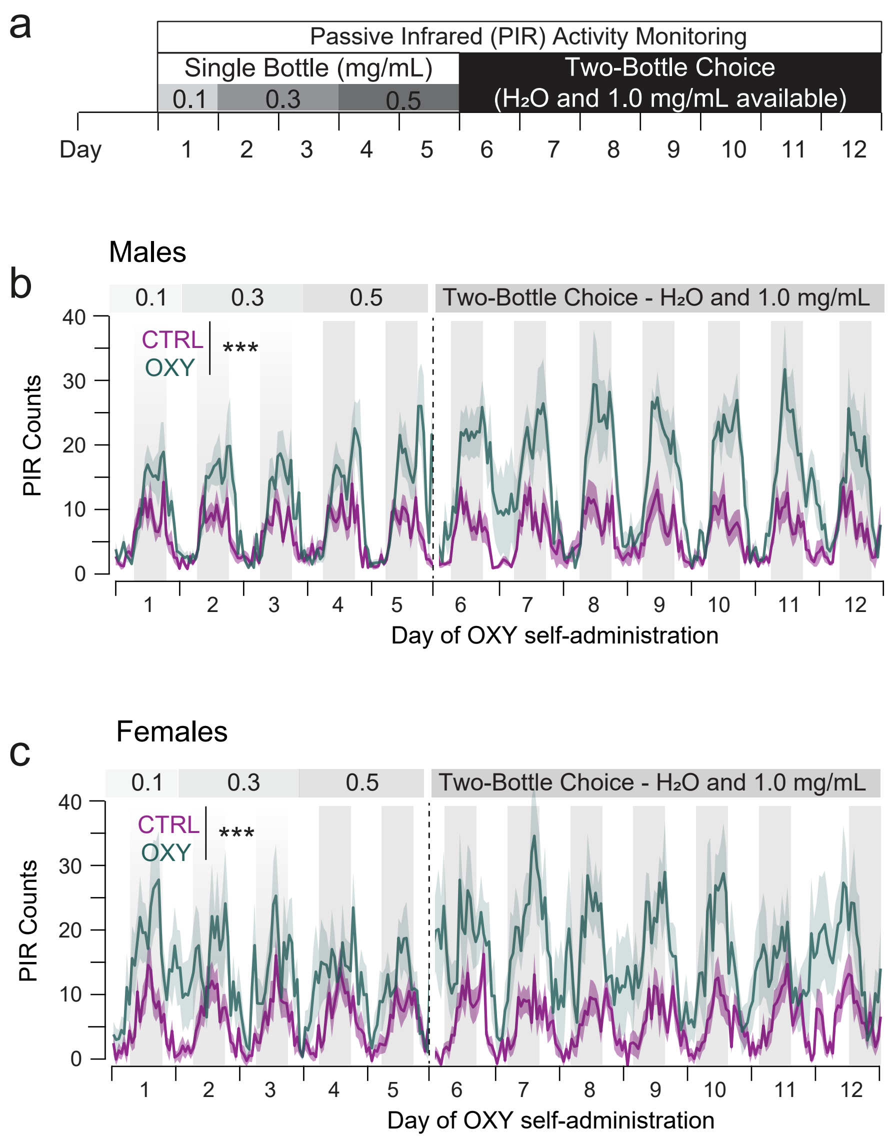


**Supplemental Figure 4. Both Male and female mice exhibited increased locomotor activity during oxycodone self-administration.**

**Supplemental Figure 4. Both male and female mice exhibited increased locomotor activity during OXY self-administration. (a)** Experimental schematic; PIR sensors monitored individual mice (CTRL: 10F/10M, OXY: 15F/13M) throughout the OXY self-administration paradigm. Increased activity was observed in both male **(b)** and female **(c)** mice. ***$p < 0.001$

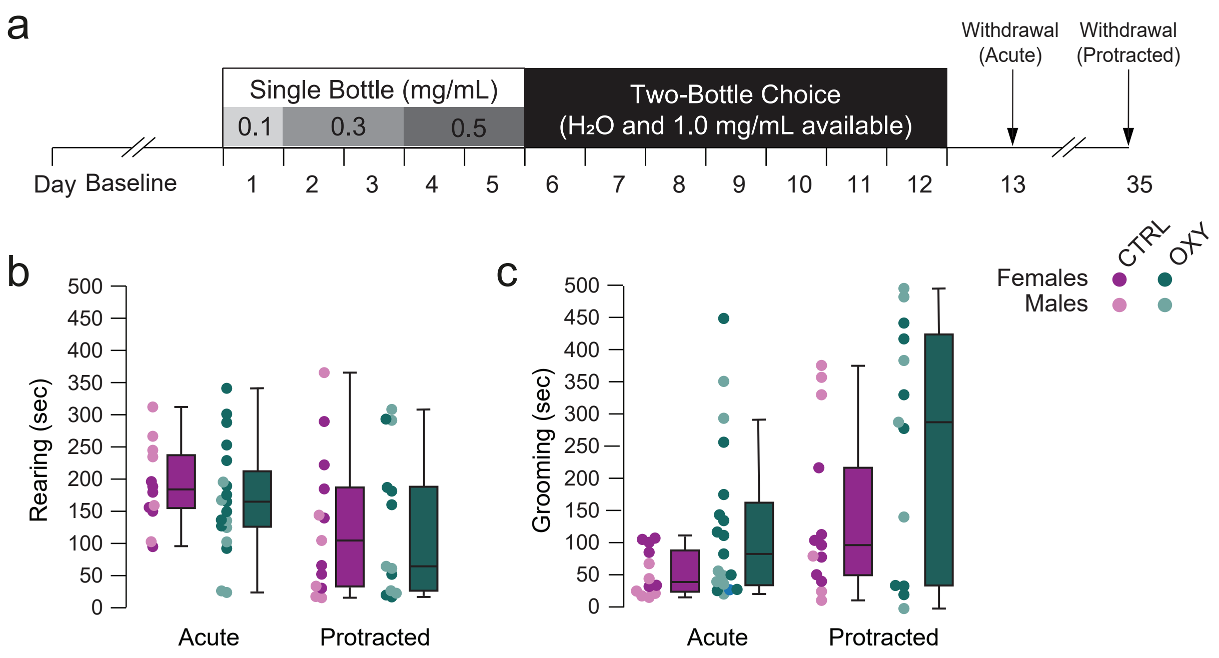


**Supplemental Figure 5. OXY self-administration has no effect on rearing or grooming behaviors in acute or protracted withdrawal.**

**Supplemental Figure 5. OXY self-administration has no effect on rearing or grooming behaviors in acute or protracted withdrawal. (a)** Experimental schematic **(b)** There was no significant effect of OXY on rearing time at either the early (CTRL: 182.39 ± 18.54, OXY: 159.66 ± 21.36) and protracted time points in either male or female mice (CTRL: 105.05 ± 22.24, OXY: 93.99 ± 23.00; $F_{Drug} = 1.359$, $p = 0.249$, $F_{Sex} = 2.011$, $p = 0.162$). **(c)** There was no significant effect of OXY on grooming time at either the early (CTRL: 59.87 ± 10.47, OXY: 118.82 ± 24.84) and protracted time points in either male or female mice (CTRL: 110.96 ± 20.29, OXY: 148.80 ± 27.94; $F_{Drug} = 3.426$, $p = 0.070$, $F_{Sex} = 0.271$, $p = 0.605$).

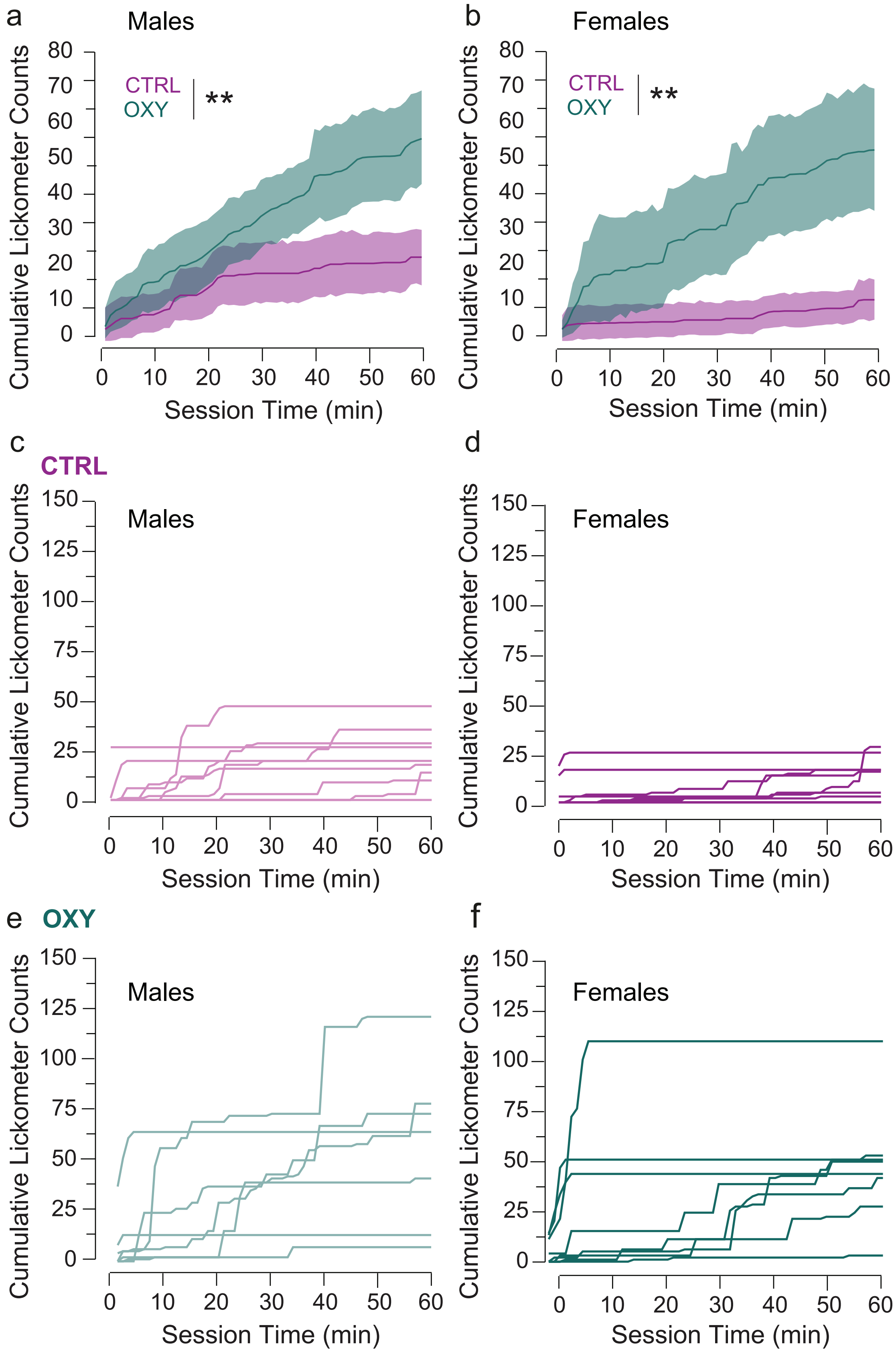


**Supplemental Figure 6. Male and female mice did not significantly differ on OXY-seeking in extinction.**

**Supplemental Figure 6. Male and female mice did not significantly differ in OXY-seeking behavior in extinction. (a-b)** Mean cumulative lickometer counts over the 60 minute probe seeking session; both male (CTRL: 20.27 ± 3.76, OXY: 55.67 ± 7.34, $t = 4.29$, $p = 0.0002$) and female (CTRL: 13.47 ± 3.011, OXY: 74.50 ± 19.34, $t = 3.19$, $p = 0.003$) OXY mice exhibited higher lickometer seeking counts relative to controls. **(c-f)** Individual lickometer seeking counts for control **(c-d)** and OXY **(e-f)** mice are shown. **$p < 0.01$

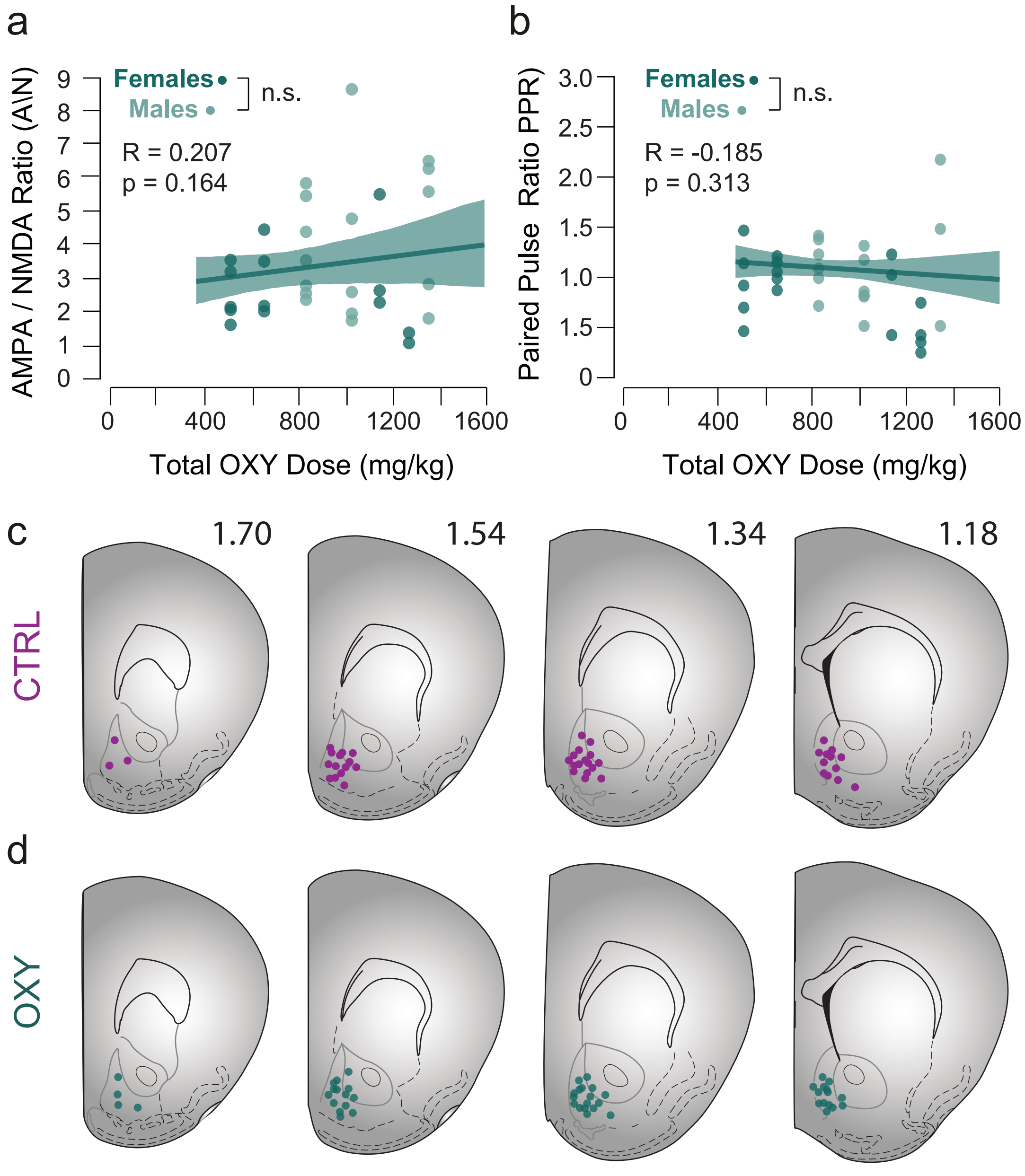


**Supplemental Figure 7. OXY intake and locations of neurons recorded with patch clamp electrophysiology.**

**Supplemental Figure 7. Locations of medium spiny neurons recorded with patch clamp electrophysiology.** Location of recorded cells in control **(a)** and OXY-self-administering groups **(b)** as registered to the Atlas of Paxinos and Watson after biocytin filling and reconstruction. AP coordinates relative to bregma are indicated (mm).
